# Dissecting specialized metabolism in space: A MALDI-MSI atlas of Amaryllidaceae alkaloids in *Hippeastrum papilio* (Ravenna) Van Scheepen

**DOI:** 10.1101/2025.07.03.663014

**Authors:** Nuwan Sameera Liyanage, Natacha Mérindol, Kenneth Munk Pedersen, Christian Janfelt, Isabel Desgagné-Penix

## Abstract

Amaryllidoideae produce specific specialized metabolites known as Amaryllidaceae alkaloids (AAs), extensively studied for their significant pharmacological potential. AAs’ spatial distribution and biosynthesis within plant tissues remain poorly understood. This study investigates organ- and tissue-specific localization in *Hippeastrum papilio*, from precursors to galanthamine and haemanthamine, using matrix-assisted laser desorption/ionization mass spectrometry imaging. Consistent accumulation of AAs was observed in epidermal and vascular tissues, with leaves exhibiting a uniform distribution across all ages and positions. Bulbs exhibited higher concentrations in the outer-scales and basal-plates, while roots displayed compartmentalized patterns, with galanthamine being uniquely abundant in the vascular bundles. Haemanthamine and galanthamine were detected in high quantities in the leaves’ and bulbs’ mucilage, while precursors were scarce. Multivariate analyses revealed that precursors clustered separately from end-products and were specifically enriched in the middle-scales and apical-leaves of the bulbs. Nonetheless, biosynthetic intermediates were observed in all tissues, indicating widespread AA biosynthesis across all organs. These findings suggest a coordinated metabolic network in *H*. *papilio*, which challenges existing hypotheses on organ-specific AA biosynthesis and hints at the transport of end-products. This study refines current models of alkaloid biosynthesis and underscore the value of *H. papilio* as a promising resource for sustainable production of therapeutic AAs.

**Highlights:** MALDI-MSI reveals widespread, tissue-specific alkaloid distribution in Hippeastrum papilio, challenging organ-specific biosynthesis and implicating epidermal and vascular tissues in Amaryllidaceae alkaloid production.

## 1. Introduction

The Amaryllidaceae (*s.s.*) comprises diverse flowering plants commonly known as amaryllis or daffodils, with notable ecological, ornamental, and pharmacological significance (Berkov *et al*., 2020). Belonging to the order Asparagales, this family includes approximately 75 genera with around 900 species, primarily characterized by their bulbous growth forms, linear leaves, and showy, often fragrant flowers (Meerow and Snijman, 1998; Meerow *et al*., 2020). They exhibit a cosmopolitan distribution, with species native to various temperate, tropical, and subtropical regions across the globe. Most Amaryllidaceae are concentrated in South America and southern Africa, serving as biodiversity hotspots for the family (Meerow, 2023; Rønsted *et al*., 2012). Other notable areas of diversity include the Mediterranean basin, Southeast Asia, and parts of Australia (Meerow, 2023). These plants are typically adapted to environments ranging from arid and semi-arid conditions to more mesic habitats, including grasslands and forest understories (Meerow and Snijman, 1998). Their ability to thrive across diverse ecological niches has contributed to their widespread cultivation and naturalization beyond their native ranges (Meerow and Snijman, 1998). Species such as *Amaryllis belladonna*, *Leucojum aestivum* (snowdrops), *Lycoris radiata* (spider lilies), *Nerine*, *Narcissus* hybrids (daffodils), and *Hippeastrum* hybrids are widely grown as ornamental plants in gardens, parks, and indoor settings.

Beyond their ornamental appeal, the Amaryllidaceae family holds significant medicinal value. For centuries, traditional medicine systems in various cultures have employed these plants to treat ailments ranging from skin disorders and wounds to more complex conditions such as epilepsy and inflammation (Jin and Yao, 2019; Nair and van Staden, 2013). Ethnopharmacological practices often rely on bulbs and leaves (Desgagné-Penix, 2021; Jayawardena *et al*., 2024; Nair and van Staden, 2013; Nair and van Staden, 2020, 2023), which contain a rich array of specialized metabolites, mainly isoquinoline-derived alkaloids with a wide range of pharmacological activities, drawing considerable attention from the scientific community (Berkov *et al*., 2020; Ka *et al*., 2020). More than 700 distinct Amaryllidaceae alkaloids (AAs) have been identified, with notable representatives, including galanthamine, lycorine, and haemanthamine (Jayawardena *et al*., 2024). Galanthamine, isolated initially from *Galanthus* species, has become a therapeutic agent for Alzheimer’s disease symptoms due to its acetylcholinesterase inhibitory activity (Heinrich and Teoh, 2004; Loy and Schneider, 2006). Lycorine has demonstrated potent antiviral, antitumor, and anti-inflammatory effects, while haemanthamine is known for its cytotoxic properties, making it a promising candidate for cancer research (Cahlíková *et al*., 2021; Jayawardena *et al*., 2024; Zhang *et al*., 2024). The current supply of these compounds is limited, primarily sourced from the extraction of only a few Amaryllidaceae species, such as *L. aestivum* and *Narcissus pseudonarcissus, Lycoris radiata, and Ungernia victoris* (Berkov *et al*., 2022; de Andrade *et al*., 2011) highlighting the need to identify more highly productive species and understand their biosynthetic pathways.

Some *Hippeastrum* species have been reported to produce high amounts of galanthamine and haemanthamine, as well as several other AAs (Figure 1), holding promise for sustainable and efficient extraction relative to other plant sources (de Andrade *et al*., 2011; Guo *et al*., 2016; Haist *et al*., 2024; Tallini *et al*., 2021). This diverse genus comprises over 100 bulbous perennial species native to South America’s tropical and subtropical regions (García *et al*., 2019). *Hippeastrum papilio*, commonly known as the Butterfly Amaryllis, stands out for its unique floral morphology and the significance of its natural habitats (Govaerts). Originating from the Atlantic forests of southern Brazil, *H*. *papilio* naturally grows on tall trees as an epiphyte, unlike most of the geophytic Amaryllidaceae species (Campos-Rocha *et al*., 2023). It is valued in horticulture for its distinctive green and burgundy flowers, often used in breeding programs to develop hybrids with enhanced ornamental traits (de Andrade *et al*., 2011; Fellers, 1998).

**Figure 1.**
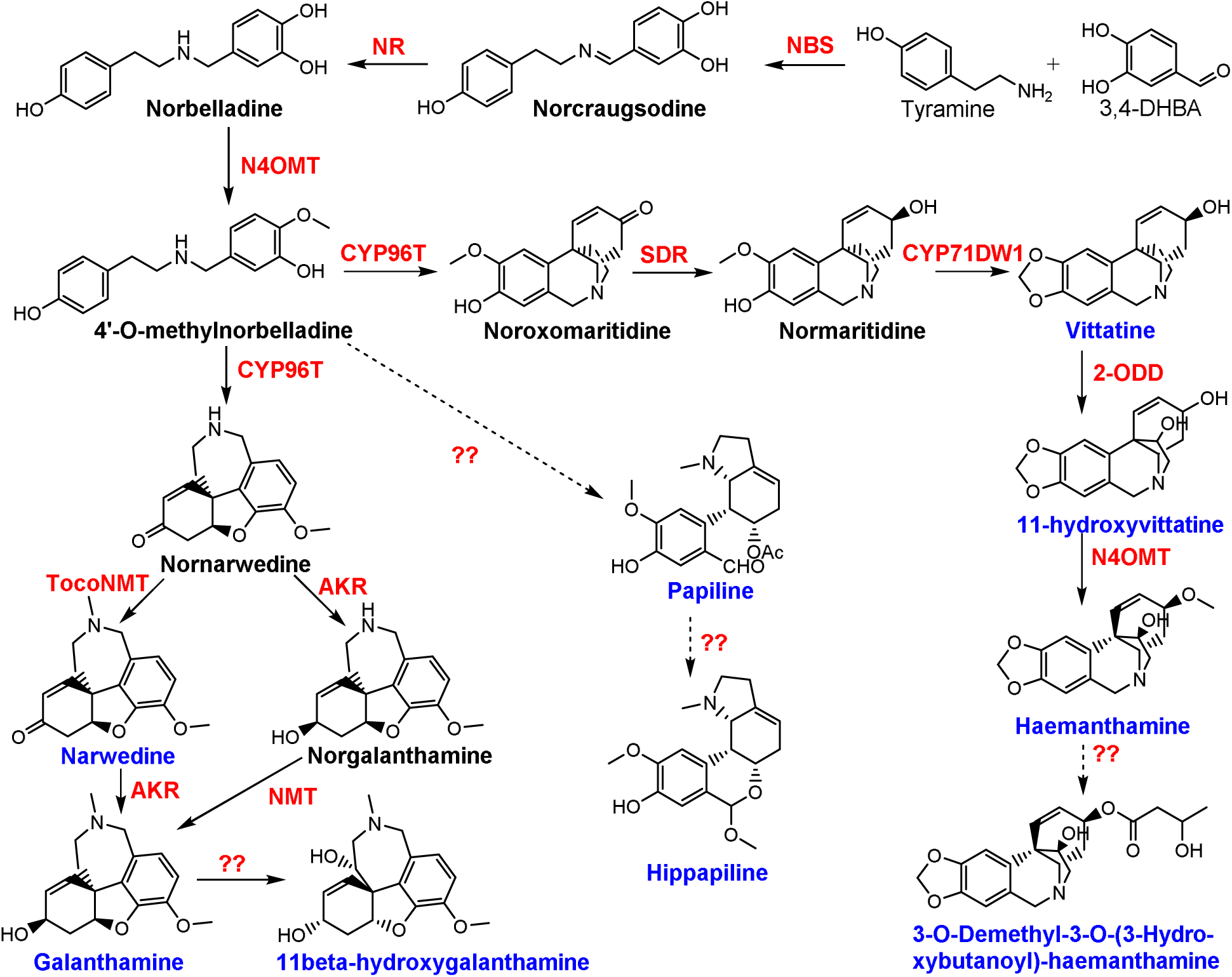
Biosynthetic pathway representing Amaryllidaceae alkaloids from *Hippeastrum papilio* starting from the initial precursors, tyramine and 3,4-dihydroxybenzaldehyde (3,4-DHBA). The bold letters (black and blue) represent the compounds examined in this study, and the blue letters indicate the alkaloids previously recorded in *H*. *papilio*. Single arrows represent a single enzymatic reaction, whereas the dotted arrows signify multiple enzymatic steps. Red letters indicate the characterized enzymatic reactions of the pathway. NBS: Norbelladine synthase, NR: Norcraugsodine/noroxomaritidine reductase, N4OMT: Norbelladine 4’-O-methyltransferase, CYP96T: Cytochrome P450 96T, TocoNMT: Tocopherol N-methyltransferase, AKR: Aldoketoreductase, SDR: Short chain dehydrogenase/reductase, NMT: N-methyltransferase, 2-ODD: 2-oxoglutarate-dependent dioxygenase (Haist *et al*., 2024; Lamichhane *et al*., 2025; Liyanage *et al*., 2025; Mehta *et al*., 2024).

Unravelling the precise tissue and organ distribution of specific alkaloids in this species is required to provide the critical knowledge to optimize the species’ industrial and pharmaceutical applications. Various techniques can be employed for the *in situ* detection of metabolites in plants, including autofluorescence, staining, labelling, and, more recently, mass spectrometry imaging (MSI) (Donaldson, 2020; Qin *et al*., 2018). Recent research has begun to explore alkaloid distribution at both histochemical and organ-specific levels of *H*. *papilio*, shedding light on their biosynthetic pathway using Dragendorff’s reagent (Haist *et al*., 2024). Haist et al. demonstrated that alkaloids are present in all plant tissues, with higher concentrations accumulating in bulbs (inner part followed by outer scales) and roots compared to leaves. Alkaloids were enriched in phloem sap of leaves, bulbs, roots, leaf parenchyma, and vascular bundles. However, Dragendorff’s staining lacks the resolution to identify individual alkaloid components and is associated with a false positivity of non-nitrogenous oxygenated compounds (Habib, 1980).

MSI combines the spatial resolution of imaging with the molecular specificity of MS (Bjarnholt *et al*., 2014). This technology works by ionizing molecules from the surface of a sample and analyzing their mass-to-charge ratios to create molecular maps that represent the spatial localization of specific compounds (Bjarnholt *et al*., 2014; Buchberger *et al*., 2017; Granborg *et al*., 2022). MSI can detect a wide range of molecules simultaneously, making it a highly efficient tool for comprehensive analysis. The technique has been employed to unravel biosynthetic pathways, allowing researchers to trace the origin, transformation, and final deposition sites of plant metabolites (Horn and Chapman, 2024; Spengler, 2015).

Matrix-assisted laser desorption/ionization (MALDI)-MSI uses a laser to desorb and ionize analytes with an applied chemical matrix, whereas desorption electrospray ionization (DESI)-MSI employs a stream of charged droplets to sample analytes directly from the surface, making it ideal for analyzing delicate plant tissues (Bjarnholt *et al*., 2014; Horn and Chapman, 2024). Monoterpenoid indole alkaloid imaging of *Rauvolfia tetraphylla* by MALDI and DESI-MSI provided detailed information on the localization of reserpine and its precursors, such as their compartmentalization in the xylem in stem tissues (Lorensen *et al*., 2023). In *Daphniphyllum macropodum,* MALDI-MSI revealed that the distribution of alkaloids within tissues is complex, likely involving intercellular transport (Eljounaidi *et al*., 2024). Alkaloids were localized in specific regions of the petiole and stem tissues, with larger C30-carbon skeleton subtypes preferentially associated with vascular tissue (phloem), and smaller C22 subtypes more frequently localized to the epidermis and enriched in immature leaves. In Amaryllidaceae, DESI-MSI showed that the leaf bases of *Narcissus* cv. Tête à Tête preferentially accumulated vittatine synthesized from radiolabelled 4’-*O*-methylnorbelladine, whereas mid-leaf accumulated non-radiolabeled lycorine (Mehta *et al*., 2024). In *Narcissus tazetta*, MALDI-MSI revealed that galanthamine was mainly concentrated in future leaves (inner bulb), while lycorine and tazettine did not exhibit a discernible distribution pattern within the bulb. No significant differences in distribution were observed among the three alkaloids in the leaf cross-sections (Nakagawa *et al*., 2024). These studies emphasize the diversity of alkaloid repartition among species, organs, tissues, and structural types, underscoring the need for further research on high-galanthamine-producing species.

By focusing on the tissue-specific localization of alkaloids in *H*. *papilio*, this study aims to enhance our understanding of its phytochemistry and potential as a high-value bioresource. We used MALDI-MS to provide a detailed analysis of the tissue distribution of AAs and their biosynthetic partners in *H*. *papilio*. A combination of principal component analysis (PCA), t-distributed stochastic neighbor embedding (t-SNE), Spearman correlation, and hierarchical clustering analyses was employed to isolate the structure- and tissue-specific distribution patterns of *H. papilio* alkaloids, aiming to elucidate their biosynthetic compartmentalization. This study addresses two critical questions regarding the spatial distribution of AAs in *H*. *papilio*: a. In which tissues and organs do galanthamine and intermediates predominantly accumulate in high-galanthamine-producing *H*. *papilio*? and b. Do other key AAs, such as haemanthamine, exhibit similar spatial arrangements within the plant tissues? Understanding these distribution patterns will provide insights into the tissue-specific biosynthesis and accumulation of these pharmacologically significant alkaloids, potentially revealing their ecological roles and the regulation of their biosynthesis.

## 2. Materials and methods

### 2.1. Plant material

Mature bulbs (four bulbs in flowering level maturity) of *H*. *papilio* were purchased from Fluwel V.O.F., Belkmerweg 20A, 1754 GB Burgerbrug, The Netherlands (https://www.fluwel.com/en-gb/products/amaryllis-papillio). The plants were grown in garden soil mixed with perlite to enhance aeration around the roots, using plastic pots placed next to a windowsill for six months, with watering as needed. Once the plants had fully entered their vegetative stage, they were uprooted, and the soil was carefully removed and thoroughly washed with tap water before being patted dry. The plants were then sectioned systematically. For the leaves, outer, middle, and inner leaves were sampled, and from each leaf, leaf tip, leaf mid, and the leaf base were separated. From the bulbs, a few 2-3 outer scales, middle scales, and inner scales, along with the basal plate sectioned for the analysis. Roots were separated into three sections: root base, middle, and tip. To minimize ice crystal formation due to the high-water content of the plant, samples were immediately dipped in 5 mM sucrose in phosphate-buffered saline for 30 minutes after sectioning. Following this, they were washed three times with distilled water to remove excess sucrose from around the tissues. After drying them quickly with tissues, the sections were frozen on a metal sheet over dry ice.

### 2.2. Sample preparation for MALDI-MSI

Samples were cryo-sectioned using techniques adapted from those developed by Kawamoto and employed for sample embedding by Montini et al (Kawamoto, 2003; Kawamoto and Kawamoto, 2021; Montini *et al*., 2020). Briefly, an n-heptane and dry ice slurry was prepared in a Dewar jar. A 2.5 × 2.0 cm mold (SECTION-LAB Co. Ltd.) was filled with a 3% (w/v) carboxymethyl cellulose aqueous gel and partially submerged in the n-heptane dry ice slurry. The samples were inserted into the embedding medium as the edges began to freeze, after which the mold was carefully immersed in n-heptane until the gel solidified completely. Once frozen, the block was removed from the mold and stored at -80°C before being cryo-sectioned. In this case it was nine leaf cross sections per block, seven cross sections from the bulb tissues per block and six cross sections of roots per block were taken in the main experiments. A Leica CM3050S cryo-microtome (Leica Microsystems, Wetzlar, Germany) was utilized to create cryo-sections of embedded plant samples at a temperature of -20 °C. The sample was appropriately trimmed using the cryo-microtome before sectioning. Tape-assisted cryo-sectioning at 10–20 μm thickness was performed by using cryo-film (SECTION-LAB Co. Ltd., Hiroshima, Japan). After sectioning, the sections were mounted on regular glass slides with the tissue side upwards using double-sided carbon tape (Electron Microscopy Sciences, Hatfield, PA, USA). The slides with the cryo-sections were stored at -80 °C until the MALDI imaging analysis.

Cryo-sectioned samples were placed in a vacuum desiccator for ten minutes for desiccation. An Olympus BH-2 microscope with reflected light was then used to capture optical images of the plant samples. Next, an iMatrixSpray was used to apply a solution of 2,5-dihydroxybenzoic acid (DHB) (30 mg/mL in 90% MeOH) as the MALDI matrix (Stoeckli *et al*., 2014). The sprayer settings were established at a height of 80 mm, with a line distance of 1 mm, a speed of 90 mm/s, a density of 3 µL/cm², and a total of 15 cycles over a 40 x 40 mm area. Following this, MALDI-MSI data were obtained after the evaluation of crystal formation using compound light microscopy.

### 2.3. MALDI-MS analysis

A Thermo QExactive Orbitrap mass spectrometer (Thermo Scientific, Bremen, Germany), equipped with an AP-SMALDI5 ion source (TransMIT GmbH, Giessen, Germany), was utilized for MALDI mass spectrometry imaging (MSI). The instrument was operated with a scan range of *m/z* 100–600 and a mass resolving power of 140,000 at *m/z* 200. Data were collected in positive ion mode (minimum n = 2 per organ). The matrix peak of DHB (*m/z* 273.03937) was used as a lock mass for internal mass calibration, ensuring a mass accuracy of ±1 ppm. The pixel size is specified for the individual images in the figure captions. Available Amaryllidaceae alkaloid standards were tested in the MALDI-MS prior to the analysis to confirm the detection of the compounds (Supplementary Fig. A1a-A1g).

### 2.5. Data processing and image generation

The “RAW + UDP to IMZML” program (version 1.6R170; TransMIT, Bremen, Germany) was used to convert the raw data files into imzML files (Schramm *et al*., 2012). Mass images were then generated using MSiReader v1.02 (Bokhart *et al*., 2017), with a mass tolerance of 1 ppm. The color scale was adjusted to enhance the visibility of the compounds of interest. The table provides the masses of the studied compounds in their protonated and deprotonated forms. Di-hexose ([C_12_H_22_O_11_ + K]+, *m/z* 381.07937) was used as a tissue marker to outline the contours. The MS images are representative of both studies and were produced using n = 2 biological replicates. All the data generated in this study was uploaded to the Metaspace metabolite annotation data platform for mass spectrometry imaging data (https://metaspace2020.org/), and annotated with KEGG-V1, ChEBI-2018-01, HMDB-v4, and NPA-2019-08 databases (Palmer *et al*., 2017).

### 2.6. Mucilage analysis

While sampling the tissues of *H*. *papilio* leaves and bulbs, the secreted mucilage was collected from three positions on the leaves (leaf tip, mid, and base) and the bulb into Eppendorf tubes, and kept on ice. To precipitate proteins, a 1:1 ratio of methanol was added to the collected mucilage (50 μL mucilage + 50 μL methanol), followed by vortexing and centrifugation at 10,000 rpm for 15 minutes. A 10 μL supernatant was diluted in 90 μL deionized water with papaverine at 1 ppm as the internal standard before analysis by LC-MS.

### 2.7. LC-MS analysis

A high-performance liquid chromatography (HPLC) system (Agilent 1100, USA) combined with a tandem quadrupole mass spectrometer (Waters Micromass Quattro Micro, USA), equipped with an electrospray ionization (ESI) source, was utilized for the alkaloid analysis. Chromatographic separation was performed using a Phenomenex Kinetex biphenyl column (75 × 2.1 mm, 2.6 μm, 100 Å pore size) maintained at a constant temperature of 30°C. A volume of five microliters from each sample was injected onto the column, which maintained a flow rate of 0.3 mL/min. The mobile phase consisted of solvent A (Milli-Q water with 0.1% formic acid) and solvent B (methanol with 0.1% formic acid). The elution gradient was as follows: from 0 to 5 minutes, isocratic at 10% B; from 5 to 7 minutes, a linear increase to 25% B; from 7 to 10 minutes, a ramp to 98% B; from 10 to 15 minutes, held at 98% B; from 15 to 16 minutes, a return to 10% B; and reconditioning from 16 to 25 minutes.

The mass spectrometer was operated under the following source conditions: a cone voltage of 30 V, a source temperature of 120°C, a desolvation temperature of 300°C, and a desolvation gas flow rate of 500 L/h. Analysis was conducted in multiple reaction monitoring (MRM) mode, with three ion transitions monitored for each compound to ensure specificity and sensitivity. Detailed information on all transitions and compound-specific parameters can be found in Supplementary Table A1. Relative quantitative analysis was conducted using calibration curves generated from standard solutions of the target alkaloids, which were processed with internal standards (compound/papaverine ratio). Data acquisition and processing were managed using MassLynx software (Waters, version details if applicable). The results were visualized and statistically analyzed using GraphPad Prism v10.

### 2.8. Dimension reduction and multivariate clustering analyses

Organ-level normalized abundance data and binary fine-tissue × organ-zone localization profiles were compiled into structured matrices for multivariate analysis. Principal component analysis (PCA) was first applied to the organ-level abundance matrix, after z-score standardization, to capture global variance in metabolite distribution patterns. A hybrid matrix was then constructed by integrating the binary fine-tissue localization data with normalized organ-level abundance values. This matrix was likewise z-score standardized and subjected to PCA to assess spatial localization and tissue-level accumulation trends jointly. t-Distributed Stochastic Neighbor Embedding (t-SNE) was performed on the hybrid matrix using the following parameters: perplexity = 5, learning rate = 200, number of iterations = 3000, and random seed = 42. This allowed for the visualization of metabolite relationships in a two-dimensional space while preserving the local neighborhood structure from the high-dimensional input. Pairwise Spearman’s rank correlation coefficients were calculated based on the hybrid matrix, and a hierarchical clustering heatmap was generated to validate metabolite groupings according to spatial and quantitative similarity. The PCA and t-SNE axes are unitless and optimized, respectively, for variance explanation and neighborhood preservation. Assistance from generative AI tools (ChatGPT, OpenAI GPT-4, 2024 release) was used in the development of the initial Python scripts used. All code was critically reviewed, iteratively adjusted, and executed locally on laboratory computers by the authors. The final analyses and figure generation were performed entirely offline, using these customized scripts in interaction with the AI to improve reproducibility, visualization clarity, and parameter selection.

Statistical differences in individual feature distributions between precursor metabolites and other alkaloid groups were assessed using a two-tailed Mann–Whitney *U* test. Hierarchical clustering was applied to the z-score standardized hybrid matrix using Ward’s linkage and Euclidean distance. Although the original matrix includes binary tissue localization features, standardization produced a continuous distribution of feature values, allowing the application of Ward’s method (Chavent *et al*., 2012; Kaufman and Rousseeuw, 2009). This approach yielded a dendrogram structure that was consistent with known biosynthetic relationships among alkaloids.

## 3. Results

The spatial distribution of AAs in-key plant structures, such as the roots, leaves, and bulbs, was analyzed using MALDI-MSI. Images of di-hexose were included in each MSI experiment, outlining the morphology and contours of the analyzed tissues and serving as internal references for spatial orientation. Compound detection was based on accurate mass measurements and comparison to authenticated standards.

### 3.1 Amaryllidaceae alkaloids in *H*. *papilio* leaf tissues

For the MALDI-MSI analysis of *H*. *papilio* leaf tissues, cross sections were prepared from three distinct regions: the innermost, middle leaf, and outermost (Figure 2a). Each leaf was further sectioned into three anatomical sections—the leaf base, mid-leaf, and leaf tip—resulting in a total of nine leaf cross sections per plant. These sections were embedded together in a 3% (w/v) carboxymethyl cellulose gel as a single block and scanned collectively to generate a comprehensive image, allowing for the comparison of signal intensities across the individual samples in the image. The leaves of *H*. *papilio* exhibited typical monocotyledonous anatomical features.

**Figure 2.**
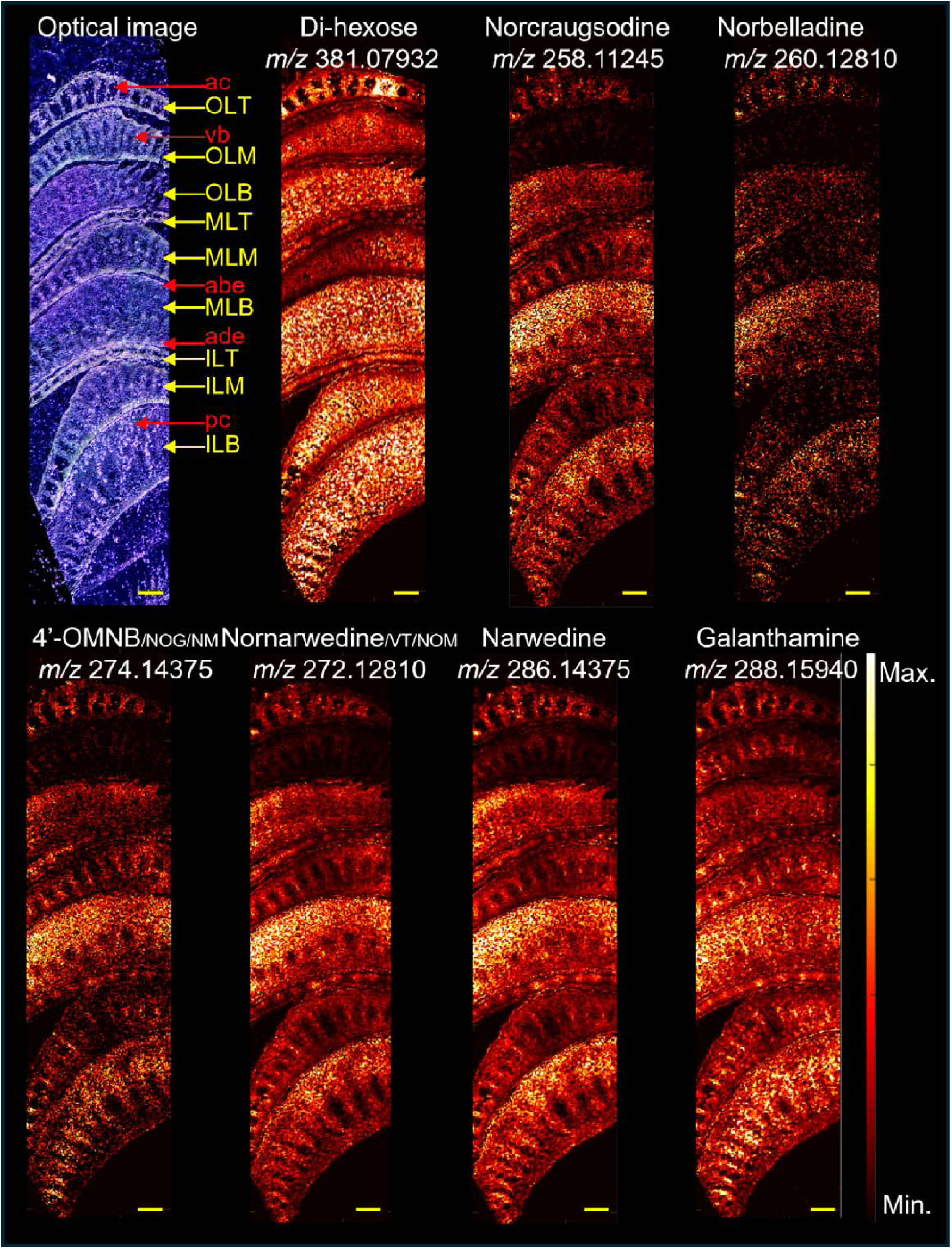
a. MALDI-MSI of *Hippeastrum papilio* leaf cross sections. From top to bottom, yellow arrows indicate the position of the leaf sections: outermost leaf (OLT = tip, OLM = mid, OLB = base), middle leaf (MLT = tip, MLM = mid, MLB = base), and innermost leaf (ILT = tip, ILM = mid, ILB = base). Red arrows indicate tissue types: ac = aerenchyma, vb = vascular bundles, abe = abaxial (lower) epidermis, ade = adaxial (upper) epidermis, and pc = parenchyma. Compound abbreviations: 4′OMNB = 4′-O-methylnorbelladine, NOG = norgalanthamine, NM = normaritidine, VT = vittatine, and NOM = noroxomaritidine. Pixel size: 40 μm. Scale bars: 1 mm.

**Figure 2.**
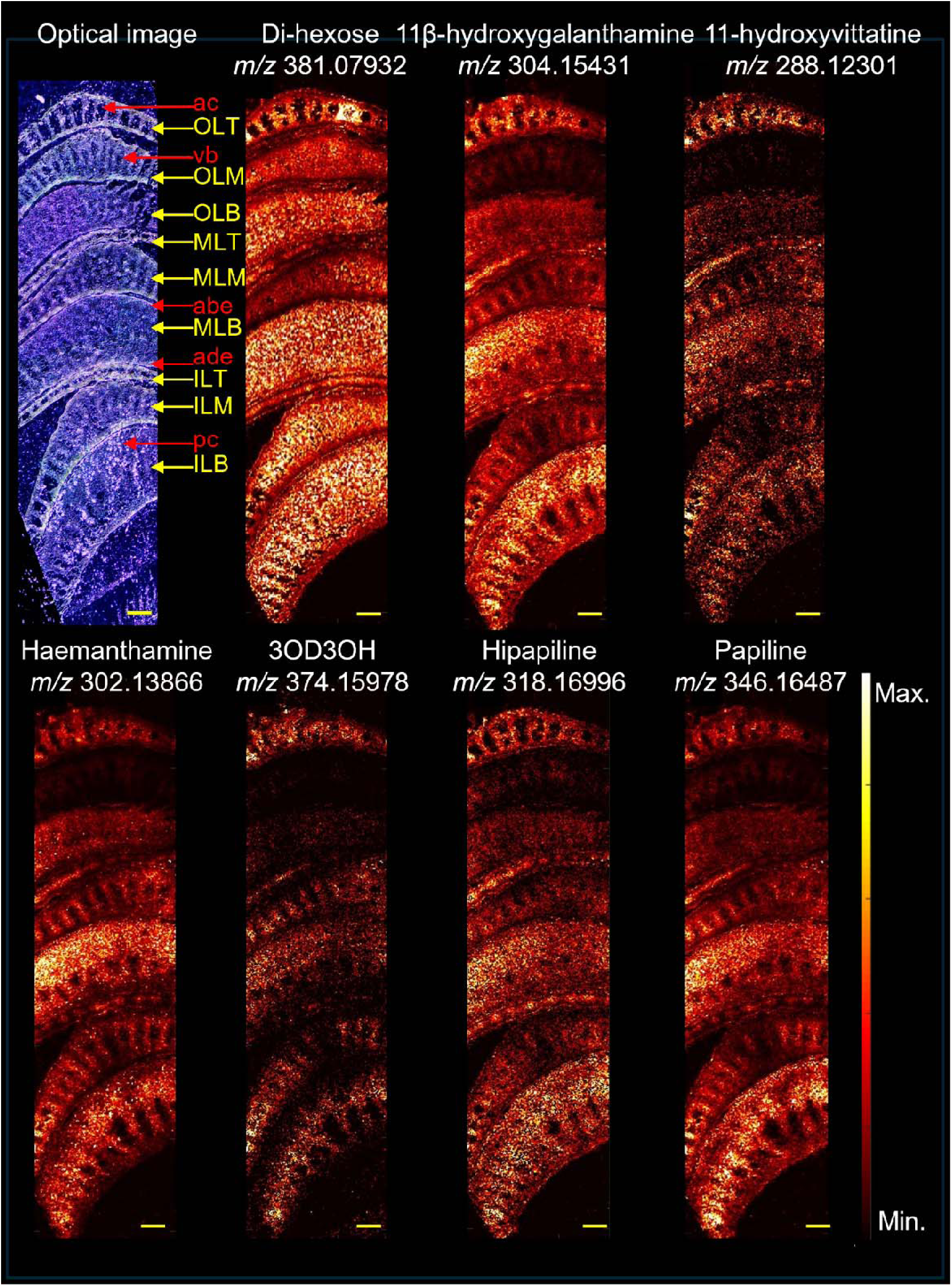
b. MALDI-MSI of *Hippeastrum papilio* leaf cross sections. From top to bottom, yellow arrows indicate the position of the leaf sections: outermost leaf (OLT = tip, OLM = mid, OLB = base), middle leaf (MLT = tip, MLM = mid, MLB = base), and innermost leaf (ILT = tip, ILM = mid, ILB = base). Red arrows indicate tissue types: ac = aerenchyma, vb = vascular bundles, abe = abaxial (lower) epidermis, ade = adaxial (upper) epidermis, and pc = parenchyma. Compound abbreviation: 3OD3OH = 3-O-demethyl-3-O-(3-hydroxybutanoyl)-haemanthamine. The pixel size is 40 μm—scale bars: 1 mm.

The MALDI-MSI focuses on the spatial distribution twelve alkaloids spanning the AA biosynthetic pathway (Figure 1), including early precursors such as norcraugsodine (*m/z* 258.1125) and norbelladine (*m/z* 260.1281) (Figure 2a; top panels), major products like galanthamine (*m/z* 288.1594) and haemanthamine (*m/z* 302.1386), and several derivatives such as 11β-hydroxygalanthamine (*m/z* 304.1543) and 3-O-demethyl-3-O-(3-hydroxybutanoyl)-haemanthamine (*m/z* 374.1597) (Figure 2a, 2b; Supplementary Table A1). Imaging was performed at a pixel size of 40 μm, with select tissues imaged at higher resolutions of 10 μm and 5 μm for finer structural detail (Supplementary Fig. A2 and in Metaspace https://metaspace2020.org/).

The spatial distribution patterns of each alkaloid were remarkably consistent across different leaf ages (outermost to innermost leaves) and positions (base, mid, and tip). Most targeted alkaloids were detected across multiple leaf regions and developmental stages, though some were absent from specific zones, such as the midsection of the outermost leaves (Figure 2a, 2b; Supplementary Table A1). Although MALDI-MSI is inherently semi-quantitative and ion intensities do not directly reflect absolute compound abundance due to potential variability in ionization efficiency, we carefully interpreted relative abundance patterns within this dataset. Notably, the alkaloids analyzed here share highly similar core structures, functional groups, and molecular weights, which likely result in comparable ionization behaviors under the applied matrix and instrumental conditions. Therefore, while we recognize that differences in ionization efficiencies may still influence signal strength, the structural similarity of the compounds provides a reasonable basis for cautious comparison of ion intensities across alkaloids in this study. This approach allowed us to gain insight into the relative distribution and accumulation trends of major versus minor alkaloids across tissues.

Among the detected signals, galanthamine and haemanthamine exhibited the highest ion intensities, which is consistent with their known accumulation as major AAs in *H. papilio*. Intermediate compounds such as norcraugsodine, norbelladine, and 4′-O-methylnorbelladine (including isomers norgalanthamine and normaritidine), as well as minor alkaloids like hippapiline and 3-O-demethyl-3-O-(3-hydroxybutanoyl)-haemanthamine, were detected at comparatively lower ion intensities. Due to potential differences in ionization efficiencies, intensities do not translate linearly into compound abundances but suggest that while the biosynthetic intermediates and minor alkaloids are present throughout the leaf tissues, their concentrations are substantially lower than the major alkaloids. Norcraugsodine, norbelladine, 4’-O-methylnorbelladine/norgalanthamine/normaritidine, nornarwedine/vittatine/ noroxomaritidine, 11-hydroxyvittatine, haemanthamine, and 3-O-demethyl-3-O-(3-hydroxybutanoyl)-haemanthamine were not detected in the midsection of the outermost leaves (Figure 2). Almost all metabolites were detected in epidermal tissues, vascular bundles, and mesophylls (parenchyma), with higher abundances observed in the epidermal and vascular regions, particularly in the lower epidermal areas of the leaf bases. In contrast, AAs were absent in the aerenchyma, which consists primarily of air spaces within the leaf tissues. Norbelladine and 3-O-Demethyl-3-O-(3-hydroxybutanoyl)-haemanthamine were not detected in outer epidermis layer, which may correspond to the cuticle, although this was not confirmed histologically. 4’-O-Methylnorbelladine/norgalanthamine/normaritidine, papilline-type, and galanthamine-type consistently accumulated in vascular bundles of most cross sections. 3-O-Demethyl-3-O-(3-hydroxybutanoyl)-haemanthamine was not detected in most epidermal tissues. Overall, galanthamine displayed the broadest and most uniform distribution across both tissues and leaf regions, highlighting its dominance among the alkaloids detected in *H. papilio* leaf tissues.

### 3.2 Amaryllidaceae alkaloids in *H*. *papilio* bulb tissues

To investigate alkaloid localization in bulb tissues of *H. papilio*, MALDI-MSI was performed on two outer scales, two middle scales, two apex leaves (leaf primordia) and a portion of the basal plate, all embedded together in a single imaging block. All twelve alkaloids previously identified in leaf tissues were detected, confirming their presence in the bulb (Figure 3a and 3b, Supplementary Table A1), but with markedly different distribution patterns. Variations in spatial localization were observed both between tissue layers and among different anatomical regions within the bulb. At the tissue level, alkaloids were predominantly detected in vascular bundles, epidermal tissues, and parenchyma.

**Figure 3.**
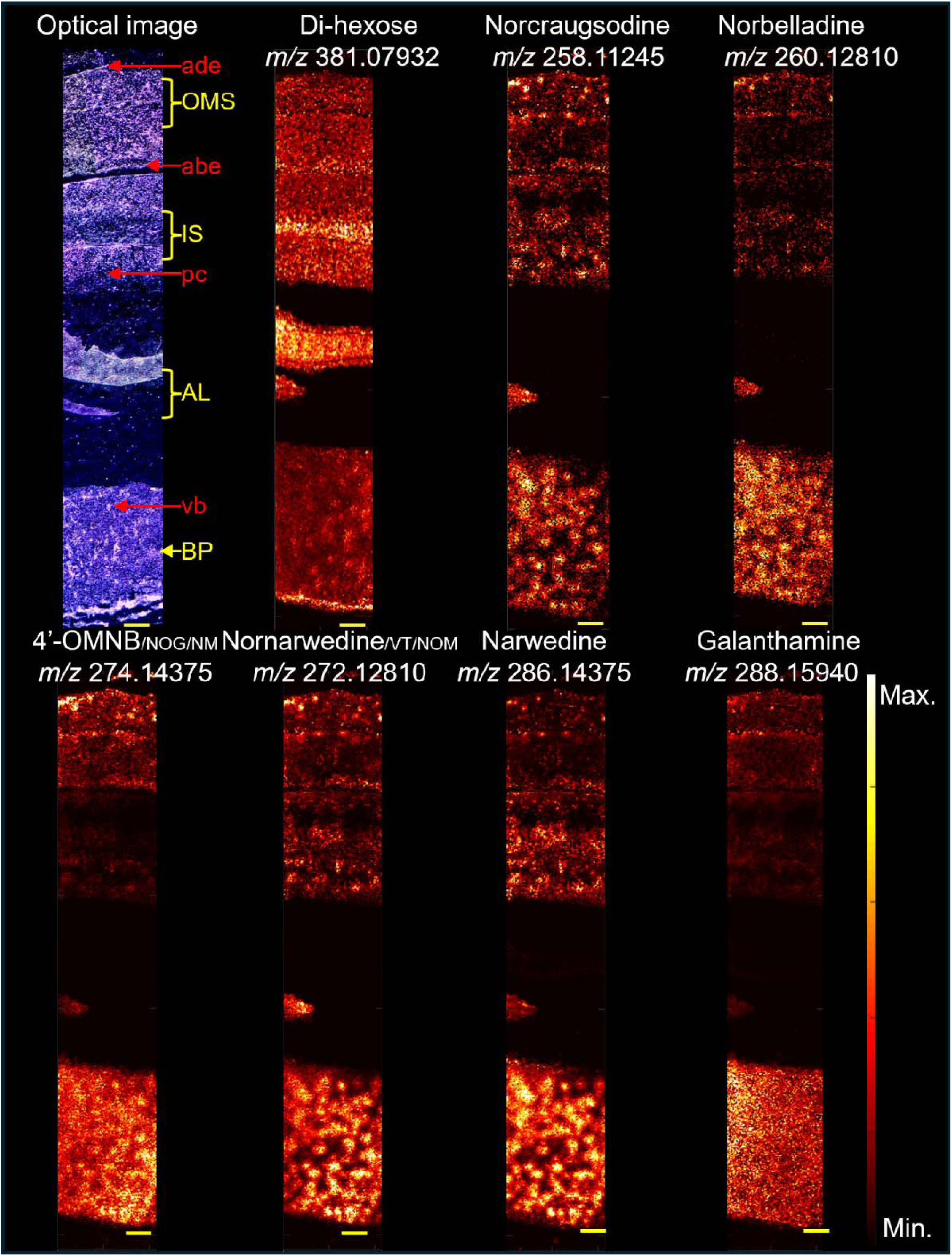
a. MALDI-MSI of *Hippeastrum papilio* bulb cross sections. In each panel, from top to bottom, yellow arrows indicate the position of the tissue sections: OMS = outermost scales (two sections), IS = inner scales, AL = apical leaves (two sections), and BP = basal plate. Red arrows denote tissue types: ade = adaxial (outer) epidermis, abe = abaxial (inner) epidermis, pc = parenchyma cells, and vb = vascular bundles. Compound abbreviations: 4′OMNB = 4′-O-methylnorbelladine, NOG = norgalanthamine, NM = normaritidine, VT = vittatine, and NOM = noroxomaritidine. The pixel size is 30 μm. Scale bars: 1 mm.

**Figure 3.**
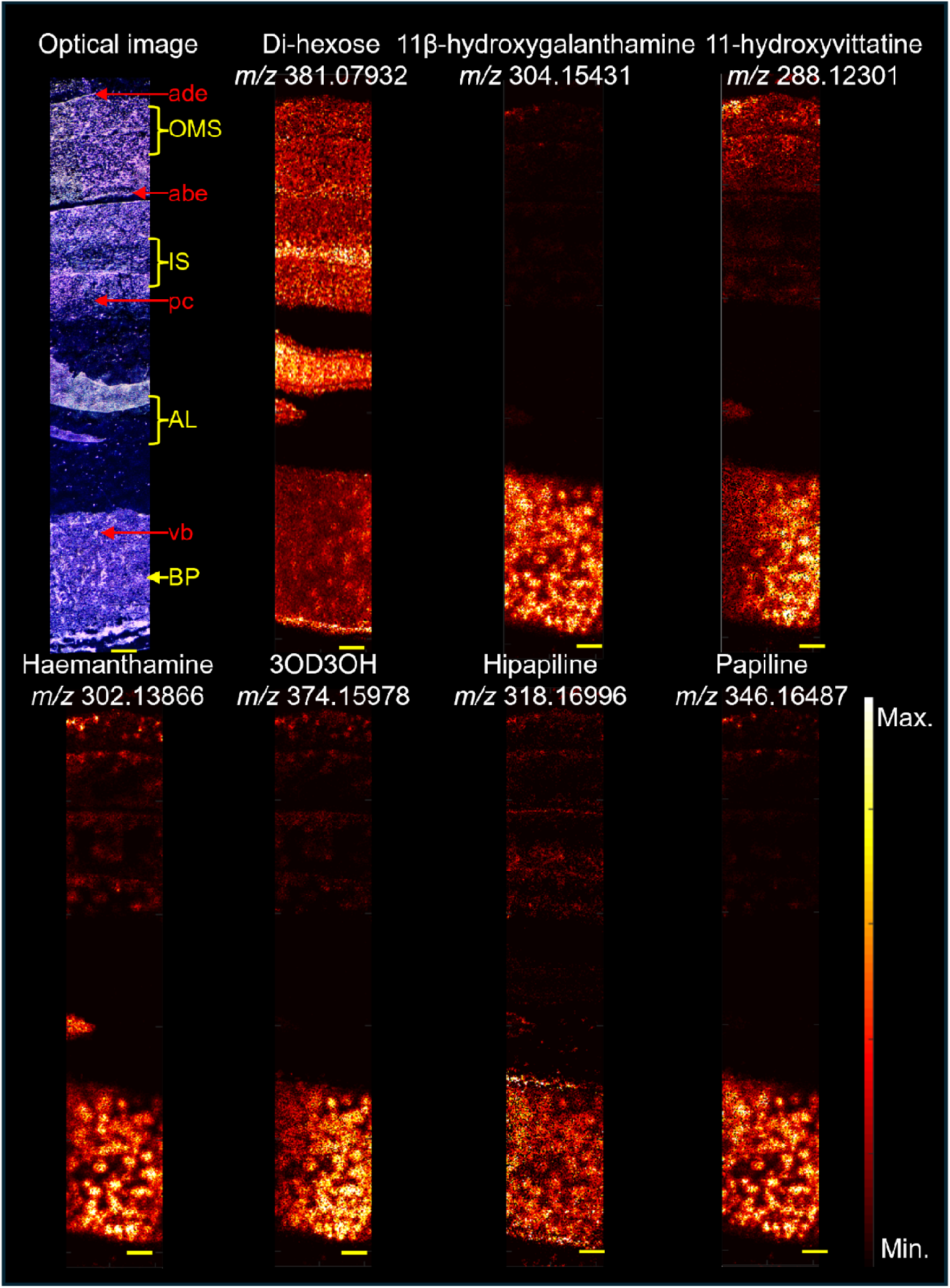
b. MALDI-MSI of *Hippeastrum papilio* bulb cross sections. In each panel, from top to bottom, yellow arrows indicate the position of the tissue sections: OMS = outermost scales (two sections), IS = inner scales, AL = apical leaves (two sections), and BP = basal plate. Red arrows denote tissue types: ade = adaxial (outer) epidermis, abe = abaxial (inner) epidermis, pc = parenchyma cells, and vb = vascular bundles. Compound abbreviation: 3OD3OH = 3-O-demethyl-3-O-(3-hydroxybutanoyl)-haemanthamine. Pixel size: 30 μm. Scale bars: 1 mm.

Alkaloid signals were primarily concentrated in the basal plate. Norcraugsodine, norbelladine, narwedine, and nornarwedine/vittatine/noroxomaritidine were consistently dispersed across both outer and inner bulb scales (Supplementary Table A1). In contrast, 4’-O-methylnorbelladine/norgalanthamine/normaritidine, galanthamine, 11-hydroxyvittatine, haemanthamine, and hippapiline were more abundant in the outer scales. Intermediates norcraugsodine, norbelladine, nornarwedine, narwedine, as well as haemanthamine, and papilline were detected in one of the apical leaves, while others were not. Norbelladine, 11β-hydroxygalanthamine, 11-hydroxyvittatine, haemanthamine, and 3-O-demethyl-3-O-(3-hydroxybutanoyl)-haemanthamine were not detected in the outer epidermis layer (possibly the cuticle) of bulb scales.

Metabolites were more abundant in the basal plates’ vascular bundles, except galanthamine, which was evenly distributed throughout the basal plate tissues. Regarding ion intensity, galanthamine, haemanthamine, 11β-hydroxygalanthamine, narwedine, and papiline were the most abundant alkaloids in *H*. *papilio* bulb tissues. These were followed by 4’-O-methylnorbelladine/norgalanthamine/normaritidine, nornarwedine/ noroxomaritidine/vittatine, and 11-hydroxyvittatine. Biosynthetic intermediates such as norcraugsodine and norbelladine, as well as minor alkaloids like hippapiline and 3-O-demethyl-3-O-(3-hydroxybutanoyl)-haemanthamine, were detected at relatively lower ion intensities. These findings suggest that while biosynthetic intermediates and minor alkaloids are present throughout the bulb tissues, their concentrations are lower compared to the major alkaloids. Overall, the distribution patterns of alkaloids in bulb tissues differed markedly from those observed in leaf tissues.

In addition to internal organ analysis, we collected mucilage from the leaf and bulb of *H*. *papilio* and measured the concentrations of alkaloids and precursors (Figure 4). Only alkaloids for which standards were available were included. Interestingly, intermediate compounds, such as norbelladine, 4’-O-methylnorbelladine, and norgalanthamine, were poorly detected in this waxy substance, except for a significant detection of 4’-O-methylnorbelladine in the mucilage coming from the bulb (119.713± 3.235 relative concentration as internal standard responsive ratio). However, galanthamine (733.26 – 1952.634 relative concentration as internal standard responsive ratio) and haemanthamine (417.205 – 1036.464 relative concentration as internal standard responsive ratio) were detected in high amounts in the mucilage of the two organs. Interestingly 4’-O-methylnorbelladine (8.11 times higher than leaf base where the highest of AA in the leaves, p = 0.0113), 11’-hydroxyvittatine (14.64 times than leaf base, p < 0.0001), galanthamine (1.57 times than leaf base, p < 0.0001) and haemanthamine (1.51 times than leaf base, p < 0.0001) were significantly higher in the mucilage extracted from the bulb compared to leaf cross-sections (Figure 4). These findings confirm that end-products accumulate in both leaves and bulbs and are present in higher amounts compared to precursors.

**Figure 4.**
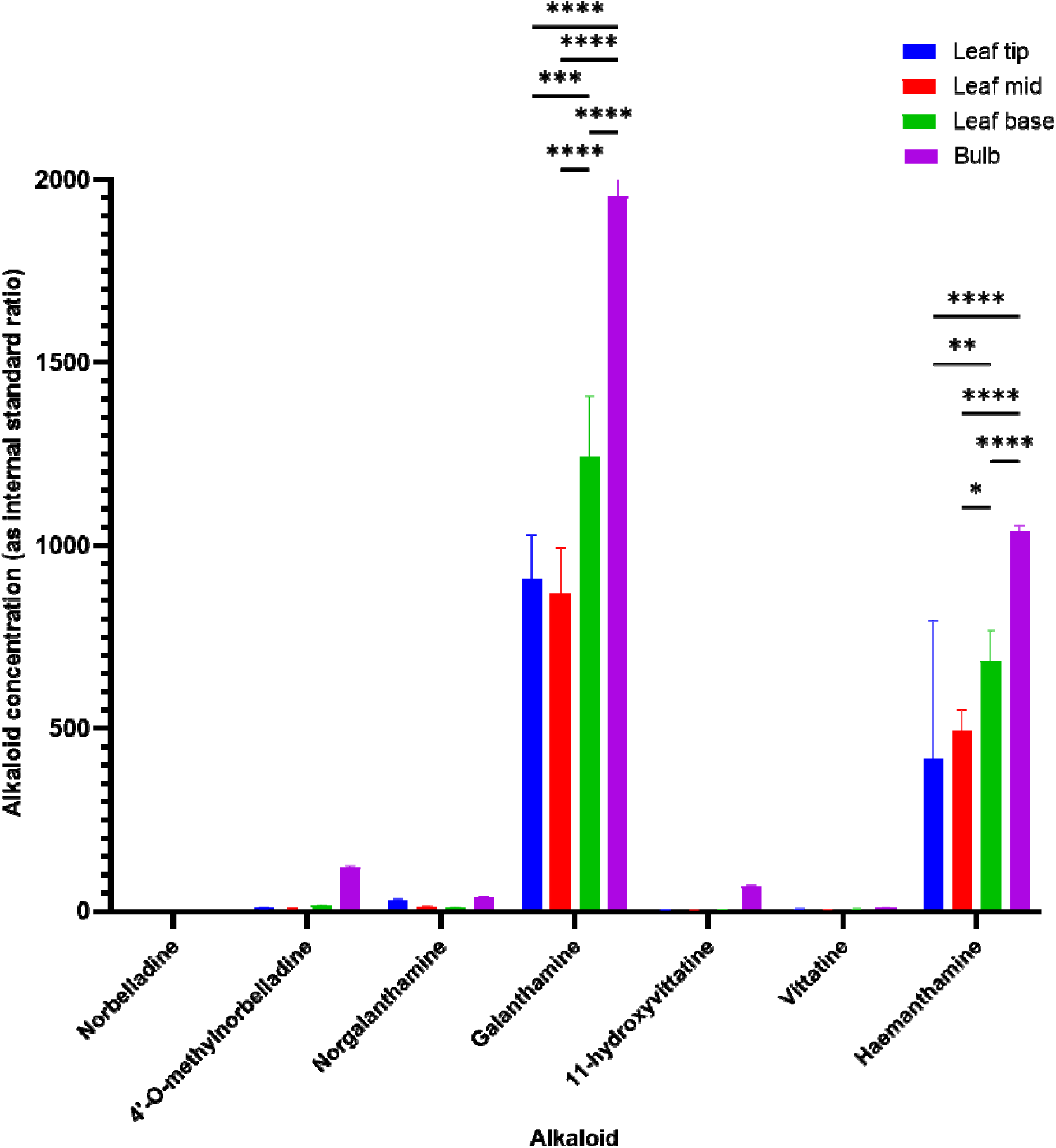
Alkaloid comparison of mucilage of *Hippeastrum papilio*. Alkaloid contents were measured b LC-MS/MS and normalized using the papaverine internal standard. All the values are shown as means ± standard deviation of three independent biological replicates. p values presented as Dunnett’s multiple comparisons test of one-way ANOVA. **** = p < 0.0001, *** = p < 0.001, ** = p < 0.01, * = p < 0.1. Non-significant interactions are not presented in the graph.

### 3.3 Amaryllidaceae alkaloids in *H*. *papilio* root tissues

To analyze the AA arrangement in the root tissues of *H*. *papilio*, single images containing two root sections from the beginning of the root, two from the middle, and two from the root tip were included to study the distribution of each AA. The MALDI-MSI analysis of alkaloids in *H*. *papilio* revealed distinct distribution patterns across root tissues compared to bulbs and leaves (Figure 5a, 5b; Supplementary Fig. A3, Supplementary Table A1).

**Figure 5.**
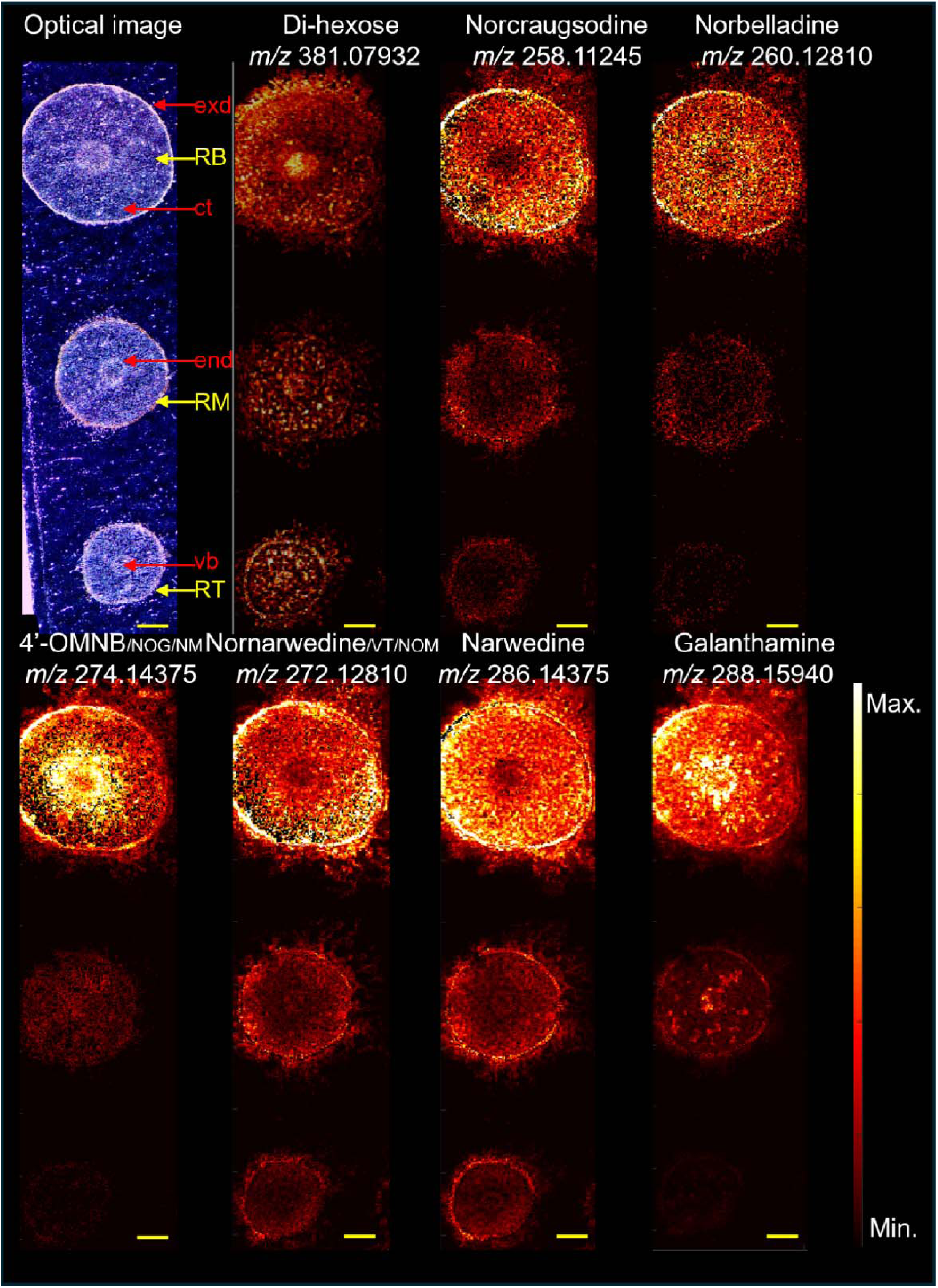
a. MALDI-MSI of *Hippeastrum papilio* root cross sections. In each panel, from top to bottom, yellow arrows indicate the position of the tissue sections: RB = root base, RM = root middle, and RT = root tip. Red arrows indicate tissue types: exd = exodermis, ct = cortex, end = endodermis, and vb = vascular bundles. Compound abbreviations: 4′OMNB = 4′-O-methylnorbelladine, NOG = norgalanthamine, NM = normaritidine, VT = vittatine, and NOM = noroxomaritidine. Pixel size: 40 μm. Scale bars: 1 mm.

**Figure 5.**
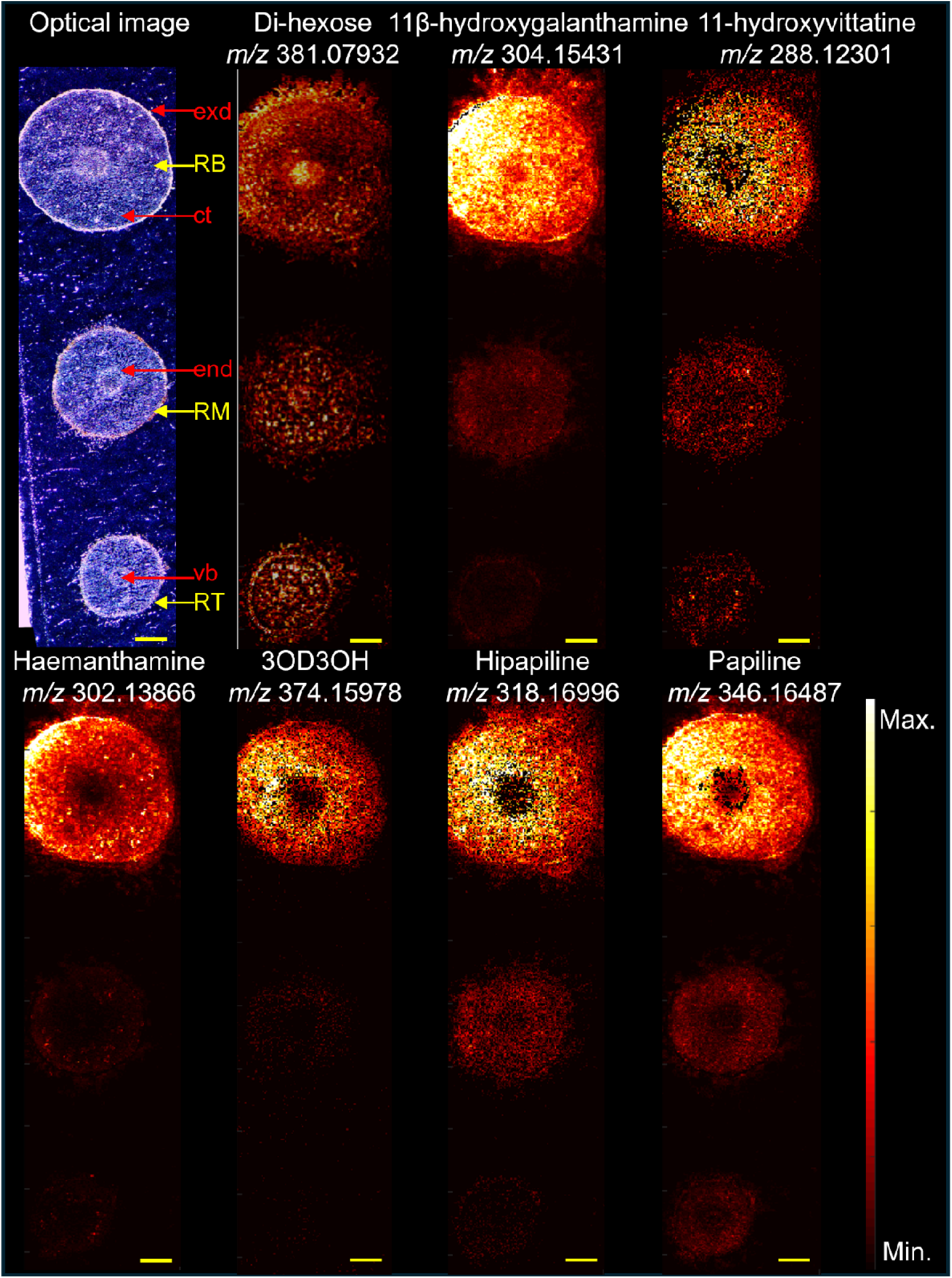
b. MALDI-MS images a cross sections of *Hippeastrum papilio* root tissues. In each panel, from top to bottom, yellow arrows indicate the position of the tissue sections: RB = root base, RM = root middle, and RT = root tip. Red arrows indicate tissue types: exd = exodermis, ct = cortex, end = endodermis, and vb = vascular bundles. Compound abbreviation: 3OD3OH = 3-O-demethyl-3-O-(3-hydroxybutanoyl)-haemanthamine. The pixel size is 40 μm. Scale bars: 1 mm.

The spatial distribution of these alkaloids varied significantly across different regions, tissue types and position along the root. At the tissue level, distinct distribution patterns emerged. Almost all tested metabolites were detected in the exodermis, cortex, and vascular bundle regions, with varying intensities (Figure 5a and 5b). For instance, norcraugsodine, norbelladine, nornarwedine/vittatine/noroxomaritidine, and haemanthamine were abundant in the exodermis and the cortex region adjacent to the exodermis (Supplementary Table A1). In contrast, 4’-*O*-methylnorbelladine/norgalanthamine/normaritidine showed higher abundance in the exodermis and cortex regions closer to the vascular bundle. Galanthamine was predominantly localized in the exodermis and vascular bundles. Hippapiline and 3-O-demethyl-3-O-(3-hydroxybutanoyl)-haemanthamine were more concentrated in the cortex, while papiline and 11β-hydroxygalanthamine were more abundant in the exodermis and cortex. Additionally, alkaloid concentrations were generally higher at the base of the roots and decreased toward the root tips. These findings suggest that the distribution of AAs in root tissues was more compartmentalized compared to leaves and bulbs.

In summary, roots of *H. papilio* accumulate a narrower spectrum of AAs compared to bulbs and leaves. The spatial distribution is predominantly restricted to the cortex and endodermis of the root base, with major alkaloids such as narwedine and papiline dominating the chemical landscape. No alkaloids were detected in the mucilage surrounding root sections (data not shown), in contrast to the alkaloid-rich mucilage of bulb and leaf tissues. The absence of early intermediates and lack of alkaloid detection in root mucilage suggest that the root may play a more limited or specialized role in alkaloid metabolism.

### 3.4 Spatial distribution and accumulation clustering of alkaloids in *H. papilio*

Hierarchical clustering revealed three primary groups that corresponded broadly to organ type—leaves, bulbs, and roots—highlighting tissue-specific accumulation profiles. Leaf and bulb samples formed two distinct but closely related clusters, reflecting their shared accumulation of major alkaloids such as galanthamine, haemanthamine, and 11β-hydroxygalanthamine, while still maintaining organ-specific differences. In contrast, root samples formed a separate and more distinct cluster, characterized by a narrower alkaloid profile and generally lower overall signal intensity (Supplementary Table A1).

To better understand patterns of localization, spatial distribution of alkaloids, and potential tissue-specific functions in *H*. *papilio*, we performed a series of multivariate analyses. A hybrid matrix was constructed that combined fine-tissue localization (presence/absence) with detection frequency across organ cross-sections. This enabled dimensionality reduction and spatial clustering of alkaloid distribution patterns. A PCA using only normalized detection frequencies across leaf, bulb, and root cross-sectional zones (tip, mid, and base) (Supplementary Figure A4) first identified metabolites based on their organ-level distribution patterns. Precursor and intermediate compounds, including norcraugsodine, norbelladine, nornarwedine, and narwedine, clustered together, reflecting similar organ cross-section-level distribution patterns. Haemanthamine was positioned close to these precursors. Most other alkaloids appeared more dispersed, indicating greater variability in their spatial profiles (Supplementary Figure A4). 4′-O-methylnorbelladine (co-detected with norgalanthamine and normaritidine) clustered in proximity to galanthamine and 11β-hydroxygalanthamine, suggesting overlapping organ-level accumulation. Conversely, 3-O-demethyl-3-O-(3-hydroxybutanoyl)-haemanthamine was positioned further from the other metabolites, reflecting a more divergent detection profile and possible tissue-specific specialization. A second PCA was performed using a hybrid matrix built from Supplementary Table A1 that combined fine-tissue × organ-zone presence/absence data with organ-level detection frequency values (Figure 6A). This integrated analysis confirmed the clear separation between precursor/intermediate metabolites (norcraugsodine, narwedine, norbelladine, nornarwedine) and more specialized derivatives, including papiline-type and galanthamine-type alkaloids. 3-O-demethyl-3-O-(3-hydroxybutanoyl)-haemanthamine remained isolated from other compounds, consistent with a distinct tissue localization profile, while 4’-O-methylnorbelladine/norgalanthamine/normaritidine position stayed close to galanthamine. Differences in tissue-level distributions, particularly in the apical bulb leaf and the tip and mid sections of the root, contributed significantly to the separation between precursor metabolites and galanthamine-type compounds (Mann–Whitney *U* test, *p* <0.05).

**Figure 6.**
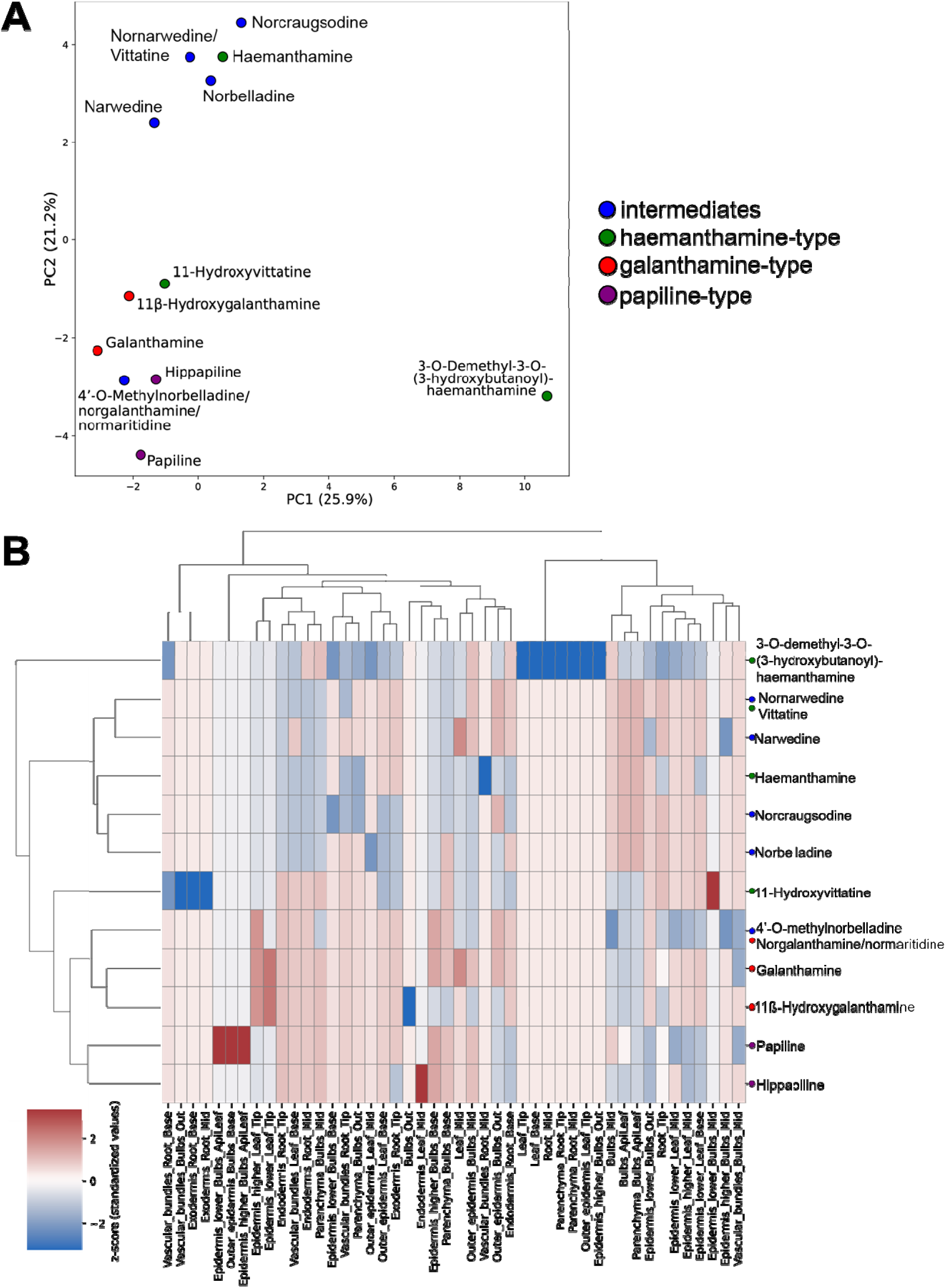
Integrated analysis of alkaloid distribution in *Hippeastrum papilio* tissues. **A.** Principal component analysis of Supplementary Table A1, integrating organ-level detection frequency and fine-tissue × organ-zone localization data. The matrix was standardized before dimensionality reduction. Metabolites are color-coded by chemical group. **B.** Hierarchical clustering of alkaloids based on their spatial distribution across organs and fine tissue zones. The hybrid matrix integrates organ-level detection frequencies and binary fine-tissue × organ-zone localization data, which are standardized using a z-score transformation. Clustering was performed using Ward’s linkage and Euclidean distance, and the resulting heatmap displays the relative spatial profiles of each alkaloid (rows) across all anatomical features (columns). Row colors denote chemical classes: blue = precursor/intermediate compounds, red = galanthamine-type, green = crinine-type, purple = papiline-type. The dendrogram reflects groupings based on overall similarity in tissue-level detection, with a cophenetic correlation coefficient r =0.77, indicating a match between clustering topology and original data structure

To verify local similarities among metabolites based on their tissue-level distribution, t-SNE was applied to the hybrid dataset (Supplementary Figure A4B). This nonlinear dimensionality reduction confirmed the distinct clustering of intermediate compounds (e.g., norbelladine, norcraugsodine, narwedine) from galanthamine-type and papiline-type alkaloids. These clusters further emphasized differences in tissue localization, particularly between compounds detected in the apical leaves of the bulb versus those enriched in the leaf tips and root tissues, reinforcing the notion of tissue-specific specialization along the biosynthetic pathway. A pairwise Spearman correlation analysis was performed on the hybrid matrix to assess similarities in metabolite distribution patterns quantitatively (Supplementary Figure A4C). Positive correlations (ρ > 0.76) were observed among intermediate compounds, consistent with their tight clustering in both PCA and t-SNE projections. Notably, narwedine and nornarwedine exhibited the highest correlation (ρ = 0.89), while galanthamine and 11β-hydroxygalanthamine also showed strong similarity in their spatial profiles (ρ = 0.85). Interestingly, across all multivariate analyses, haemanthamine consistently clustered more closely with intermediate precursors. To further explore alkaloid relationships based on tissue localization patterns, we applied hierarchical clustering using Ward’s method to the hybrid matrix (Figure 6B). This analysis grouped the twelve alkaloids into four main clusters. To assess the reliability of the hierarchical clustering structure, we computed the cophenetic correlation coefficient between the original distance matrix and the dendrogram derived from Ward’s method. The resulting coefficient r = 0.77 indicates concordance between the dendrogram topology and the underlying data structure, supporting the robustness of the observed alkaloid groupings. The first cluster included the majority of precursor and intermediate compounds—norcraugsodine, norbelladine, nornarwedine (vittatine/noroxomaritidine), and narwedine—as well as haemanthamine. These compounds were predominantly detected in the apical bulb leaf and the mid-section of the root, and showed limited presence in vascular zones. The second cluster comprised 4’-O-methylnorbelladine/norgalanthamine/normaritidine, galanthamine, 11ß-hydroxygalanthamine, and the papiline-type alkaloids (papiline and hippapiline), which were more frequently detected in vascular bundles, leaf base, and root tip tissues, including the endodermis and epidermis. A third cluster contained only 11-hydroxyvittatine, characterized by detection in both cortex and vascular regions of the root and bulb, suggesting a partially overlapping but distinct localization. The fourth cluster grouped 3-O-demethyl-3-O-(3-hydroxybutanoyl)-haemanthamine, which showed a unique pattern with detection restricted to bulb parenchyma and mid-root cortex but absent from most other tissue zones. These groupings were also consistent with the spatial separation patterns observed in the PCA and t-SNE analyses.

### 3.5. Additional compound annotations with Metaspace

In addition to using the comprehensive literature review on alkaloids reported in *H*. *papilio* as database, we utilized the MSI metabolite annotation platform Metaspace (Palmer *et al*., 2017) to analyze the data obtained from our non-targeted MSI experiments. Even if the identification in Metaspace is based solely on accurate mass analysis (leaving the possibility open for the presence of several isomeric compounds instead of or in addition to the tentatively identified compound), it revealed the putative presence of multiple classes of compounds (Supplementary Table A2), such as colchicine derivatives (deacetylcolchicine, N-acetoacetyl-deacetylcolchicine, trimethylcolchicinic acid/ N-deacetylcolchiceine, demecolchine/androcymbine/isoandrocymbine) (Supplementary Fig. A5E-H), flavonoids (kaempferol-3-O-rutinoside/ kaempferol 3-O-rhamnoside-7-O-glucoside, luteolin/kaempferol, quercetin 3-O-glucoside, quercetin/8-hydroxykaempferol/6-hydroxykaempferol/8-hydroxyluteolin. 4’-methoxyflavanone), amides (N-*p*-trans-coumaroyltyramine, N-trans-caffeoyltyramine), lignans (1-acetoxypinoresinol, deoxypodophyllotoxin), pigments (pheophorbide a, adonixanthin/ dinoxanthin/ didinoxanthin, echinenone, 4-ketomyxol), and fungicide (flumetover) (Supplementary Fig. A6, Supplementary Table A2)

Metaspace annotations also indicated the potential presence of AAs previously documented in other species of the Amaryllidaceae family but not yet reported in *H*. *papilio*. These findings are summarized in Figure 7, the accompanying Supplementary Table A2, and Supplementary Fig. A5, which provide detailed insights into the newly identified AAs and their potential biological and ecological significance. These included phenanthridine-type alkaloids such as crinasiadine and trisphaeridine, β-carboline alkaloids like trichotamine, and a variety of AA, including assoanine, oxoassoanine, albomaculine, ungeremine, hippeastrine, crinan/gamma-lycorane, O-methyllycorenine, belladine, amaryllisine/lycorenine, and papyramine. Additionally, metabolites associated with the alkaloid degradation pathway, such as galanthamine beta-D-glucuronide and O-demethyl-galanthamine beta-D-glucuronide, were identified. These findings highlight the chemical diversity of *H*. *papilio*.

**Figure 7.**
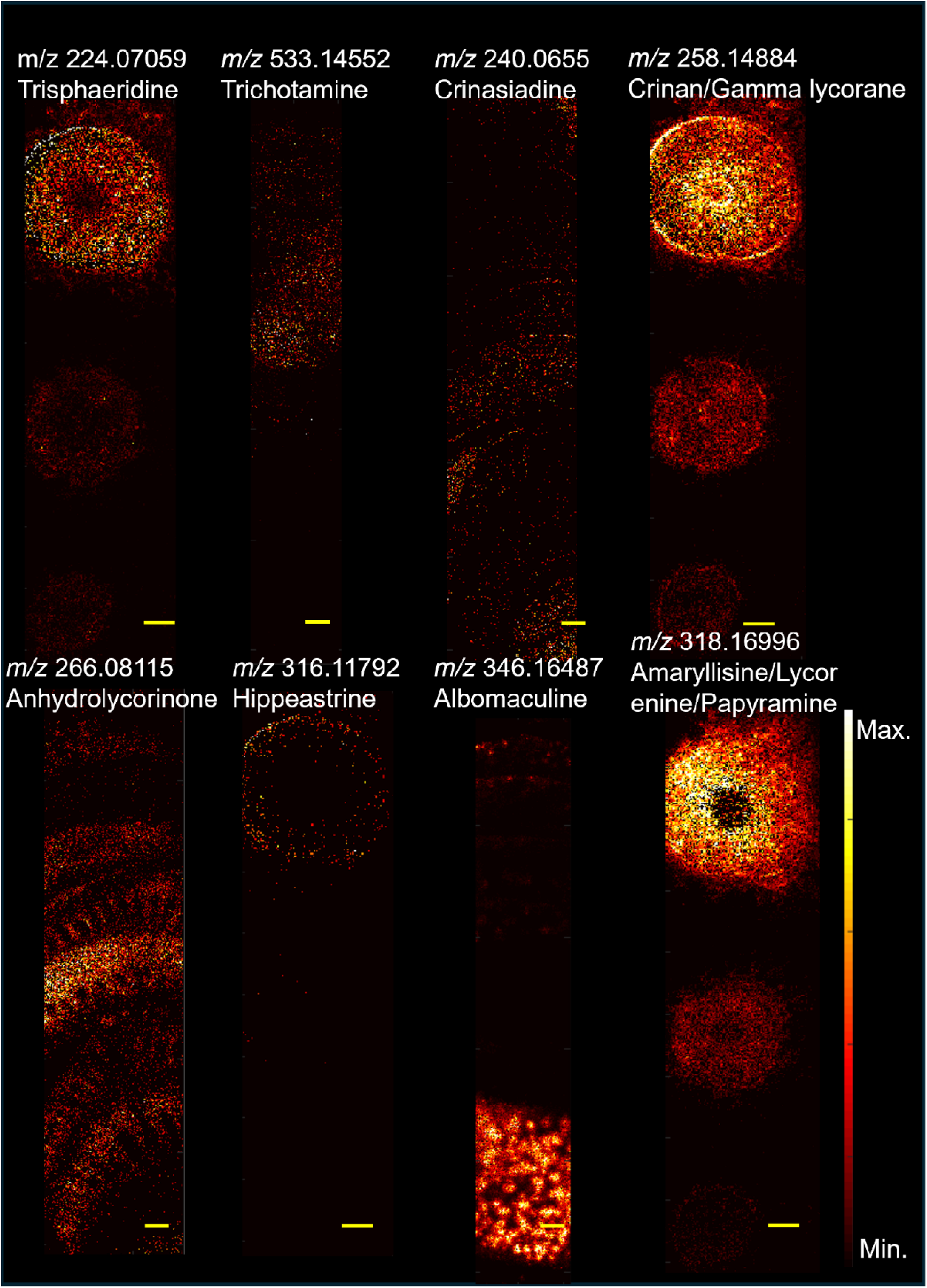
MALDI-MSI of multiple METASPACE-annotated compounds in cross sections of *Hippeastrum papilio* tissues. Panels A, D, F, and H show root sections (pixel size: 40 μm); panels B and G show bulb sections (pixel size: 30 μm); and panels C and E show leaf sections (pixel size: 40 μm). Scale bars: 1 mm.

## 4. Discussion

In this study, MALDI-MSI and LC-MS were used to gain insight into alkaloid biosynthesis in high levels of galanthamine-producing *H. papilio,* by analyzing the spatial distribution in different organs. One of the primary strengths of MSI is its label-free nature, which enables the direct analysis of a wide range of molecules simultaneously in various samples (Spengler, 2015). This feature preserves sample integrity and reduces preparation time (Dong *et al*., 2016). It has revolutionized numerous scientific fields by enabling the visualization of the spatial distribution of molecules (Buchberger *et al*., 2017). Unlike traditional MS, which provides bulk molecular information, MSI retains the spatial context of analytes, making it invaluable for understanding complex biological systems (Buchberger *et al*., 2017; Qin *et al*., 2018). Over the years, MSI has become an indispensable tool across various disciplines, from medicine and drug development to agriculture (Buchberger *et al*., 2017; Granborg *et al*., 2022), enabling researchers to map the spatial distribution of metabolites directly on plant tissues (Bjarnholt *et al*., 2014). MALDI-MSI results are inherently qualitative for this type of plant tissue analysis. While the imaging data reveals the spatial distribution of compounds, it does not provide quantitative information. Variations in ionization efficiencies among different compounds also limit the direct comparison of their concentrations. In *H. papilio*, we first focused on the repartition of well-known alkaloids end products (such as galanthamine, haemanthamine) and intermediates (such as norbelladine, norcraugsodine) in cross-sections of three organs (leaves, bulbs and roots). The presence of these specific alkaloids in each cross-section was deeper analyzed based on detection in specific tissues (e.g. parenchyma, epidermis). One limitation of the current analysis was the presence of isomers (same m/z) for 4’-O-methylnorbelladine (norgalanthamine, normaritidine) and nornarwedine (vittatine/ noroxomaritidine), which introduced signal overlap in these cases (Table 1).

**Table 1.** Exact masses and ionized forms of the compounds imaged by MALDI-MSI.

| Alkaloid | Molecular<br>formula | Monoisotopic<br>mass (Da) | [M+H]<sup>+</sup> | [M+K]<sup>+</sup> | Known isomers |
| --- | --- | --- | --- | --- | --- |
| Norcraugsodine | C<sub>15</sub>H<sub>15</sub>NO<sub>3</sub> | 257.10518 | 258.11245 | 296.06833 | n.a. |
| Norbelladine | C<sub>15</sub>H<sub>17</sub>NO<sub>3</sub> | 259.12083 | 260.12810 | 298.08398 | n.a. |
| 4'-O-Methylnorbelladine | C<sub>16</sub>H<sub>19</sub>NO<sub>3</sub> | 273.13648 | 274.14375 | 312.09963 | Norgalanthamine,<br>Normaritidine |
| Nornarwedine | C<sub>16</sub>H<sub>17</sub>NO<sub>3</sub> | 271.12083 | 272.12810 | 310.08398 | Noroxomaritidine,<br>Vittatine |
| Narwedine | C<sub>17</sub>H<sub>19</sub>NO<sub>3</sub> | 285.13648 | 286.14375 | 324.09963 | n.a. |
| Norgalanthamine | C<sub>16</sub>H<sub>19</sub>NO<sub>3</sub> | 273.13648 | 274.14375 | 312.09963 | 4'-O-methylnorbelladine,<br>Normaritidine |
| Galanthamine | C<sub>17</sub>H<sub>21</sub>NO<sub>3</sub> | 287.15213 | 288.15940 | 326.11528 | n.a. |
| 11β-Hydroxygalanthamine | C<sub>17</sub>H<sub>21</sub>NO<sub>4</sub> | 303.14704 | 304.15431 | 342.11019 | n.a. |
| Noroxomaritidine | C<sub>16</sub>H<sub>17</sub>NO<sub>3</sub> | 271.12083 | 272.12810 | 310.08398 | Nornarwedine, Vittatine |
| Normaritidine | C<sub>16</sub>H<sub>19</sub>NO<sub>3</sub> | 273.13648 | 274.14375 | 312.09963 | 4'-O-methylnorbelladine,<br>Norgalanthamine |
| Vittatine | C<sub>16</sub>H<sub>17</sub>NO<sub>3</sub> | 271.12083 | 272.12810 | 310.08398 | Nornarwedine,<br>Noroxomaritidine |
| 11-Hydroxyvittatine | C<sub>16</sub>H<sub>17</sub>NO<sub>4</sub> | 287.11574 | 288.12301 | 326.07889 | n.a. |
| Haemanthamine | C<sub>17</sub>H<sub>19</sub>NO<sub>4</sub> | 301.13139 | 302.13866 | 340.09454 | n.a. |
| 3-O-Demethyl-3-O-(3-hydroxybutanoyl)-haemanthamine | C<sub>20</sub>H<sub>23</sub>NO<sub>6</sub> | 373.15251 | 374.15978 | 412.11566 | n.a. |
| Papiline | C<sub>19</sub>H<sub>23</sub>NO<sub>5</sub> | 345.15760 | 346.16487 | 384.12075 | n.a. |
| Hippapiline | C<sub>18</sub>H<sub>23</sub>NO<sub>4</sub> | 317.16269 | 318.16996 | 356.12584 | n.a. |
| Di-hexose (Sucrose) | C<sub>12</sub>H<sub>22</sub>O<sub>11</sub> | 342.11616 | 343.12344 | 381.07932 | n.a. |

n.a.: not applicable

### 4.1. Leaf tissues

Previous studies have suggested that AAs are biosynthesized in young leaf tissues (Mehta *et al*., 2024). Consistently, all AAs were detected in the innermost part of the leaf. In contrast, intermediates like norcraugsosine, norbelladine, 4’-O-methylnorbelladine (norgalanthamine, or normaritidine), nornarwedine (vittatine/noroxomaritidine) were detected in some but not all outermost sections of the leaf. Yet, the detection of these intermediates in the base and tip leaf sections suggests that alkaloid biosynthesis could be a spatially widespread process, not confined to specific regions. Their detection at low ion intensities could indicate transient accumulation in the tissues, consistent with their role of precursors for the synthesis of more complex alkaloids (Li *et al*., 2024). The distribution of galanthamine, papiline, and hippapiline across different leaf ages and positions suggests that these compounds may play a generalized role in leaf physiology and serve analogous functions, such as protecting photosynthetic tissues, maintaining leaf structural integrity, and defending against pathogens and herbivores, rather than being specific to certain developmental stages or regions. This observation is consistent with the findings of Nakagawa et al. (2024), who reported no significant differences in the distribution patterns of galanthamine, lycorine, and tazettine in *N*. *tazetta* leaf tissues (Nakagawa *et al*., 2024).

While our study did not achieve subcellular resolution, the ultra-sensitivity of the MALDI-MSI instrumentation allowed the detection of alkaloids in specific tissues. Mehta et al. (2024) observed that alkaloids in *Narcissus* cv. Tête-à-Tête were localized near vascular tissues of leaves, a finding that resonates with the results of this study (Mehta *et al*., 2024). The abundance of alkaloids in epidermal and vascular tissues, particularly in the lower epidermal regions of the leaf bases, underscores the potential role of these compounds in defence mechanisms (Tissier *et al*., 2014). Epidermal tissues are the first living cells of a plant body that separate the plant from the environment, hence the presence of alkaloids in these regions may deter herbivory and microbial infections (Dietz and Hartung, 1996). Similarly, the localization of alkaloids in vascular bundles suggests their involvement in systemic defence and possibly their transport (Hagel *et al*., 2012). The absence of alkaloids in the aerenchyma supports the idea that these compounds are strategically localized in tissues with higher metabolic activity and defence requirements. This finding aligns with the histochemical localization study by Haist et al. (2024), which detected alkaloids in the cuticle, vascular bundles, intracellular spaces, nucleus, and vacuoles of *H*. *papilio* leaf tissues (Haist *et al*., 2024).

### 4.2. Bulb tissues

Compared to leaves, there was greater variation in AA repartition across the bulbs’ cross-sections. AA accumulated intensely in the basal plates, followed by the outermost bulb scales and the middle bulb scales. Intriguingly, while most intermediates were detected in apical leaves, end-products AAs, except haemanthamine, were not. The differential distribution suggests that they may serve distinct functional roles depending on their localization. For instance, compared to middle scales and apical leaves, the higher abundance of galanthamine in the outer bulb scales and basal plates, which are more exposed to the external environment, implies a potential defensive role against herbivores or pathogens. Our results differ from those of Haist et al., which showed a high accumulation of all AAs, including galanthamine, in the inner bulb sections of *H. papilio* using a less sensitive method (Haist *et al*., 2024). Our results cannot be directly compared either with those of Mehta et al., who excluded apical leaves from their analysis (Mehta *et al*., 2024). Most Amaryllidaceae species undergo hibernation periods during unfavourable conditions, such as winter or drought, where the bulbs are the only organs that remain. Therefore, bulbs should accumulate a high amount of defense-responsive molecules to protect the plant. Our mucilage analysis also supports this idea that the AAs in the mucilage of bulbs are higher than in leaves. Nevertheless, the accumulation profile of galanthamine does not align with the findings by Nakagawa et al. (2024), who reported higher galanthamine levels in the apex leaves of *N*. *tazetta* bulbs (Nakagawa *et al*., 2024). Their study did not examine the basal plates. Compared to the subterranean bulbs of *N*. *tazetta, H*. *papilio* is an epiphyte, and its bulb is more exposed, which may necessitate different defense strategies. The higher concentrations of AAs in epidermal and vascular tissues further support their involvement in defense mechanisms. Consistent with our analysis of the bulb, histochemical staining studies in *H*. *papilio* indicated that AAs were primarily localized in intercellular spaces, vacuoles, and nuclei, rather than in outer cell layers, such as the cuticle (Haist *et al*., 2024). The abundance of alkaloids in the basal plates, particularly around vascular regions, suggests a potential role in nutrient transport or storage. While most alkaloids were more specific to the vascular bundles, galanthamine was present in both the parenchymal and vascular bundles of the basal plates. This reinforces the particular physiological function of this metabolite, while its ubiquity is consistent with its high abundance in the species.

### 4.3. Root tissues

A unique distribution pattern of alkaloids was observed in *H*. *papilio* root tissues, exhibiting more complex and compartmentalized features compared to leaves and bulbs. One of the most intriguing observations was a specific intense accumulation of galanthamine, and to a lesser extent, of 4’O-methylnorbelladine/norgalanthamine/normaritidine and 11β-hydroxygalanthamine in the vascular regions of the roots. This contrasts with leaves and bulbs, where vascular areas showed a high abundance of most alkaloids. Previous studies, such as those on *N*. *tazzetta*, *N.* cv. Tête-à-Tête, did not investigate root tissues, preventing the possibility of comparison (Mehta *et al*., 2024; Nakagawa *et al*., 2024). Histochemical staining of *H*. *papilio* roots has previously indicated that alkaloids are present in all three main regions—exodermis, cortex, and vascular cylinder, with higher intensity in the vascular area (Haist *et al*., 2024). However, these findings represent the cumulative presence of all AAs, making it difficult to compare with the specific distribution patterns of individual alkaloids observed in this study. As epidermal and exodermal tissues are exposed to the environment, the chemical defense mechanism might be highly active in that area. For example, it has been shown that defense-responsive genes are expressed in epidermal cells of wheat in response to pathogen attacks (Altpeter *et al*., 2005). The compartmentalized distribution of AAs in root tissues suggests a highly regulated biosynthetic pathway, which may be tailored to the specific physiological and ecological roles of these compounds in *H*. *papilio*.

### 4.4 Differences in the precursors’ distribution in *H. papilio*

Multivariate analysis in MSI studies often relies on quantitative ion intensities. Instead, we applied a hybrid matrix integrating binary fine localization patterns and the normalized frequency of organ-level detection. This structured approach aimed to enhance the biological interpretation of spatial metabolite compartmentalization. PCA, t-SNE, Spearman, and hierarchical clustering all revealed a clear spatial separation between precursor alkaloids (e.g., norbelladine, norcraugsodine, narwedine, nornarwedine/vittatine/noroxomaritidine) and downstream galanthamine-type and papiline-type compounds, with haemanthamine clustering with precursors. This observation suggests that early and late stages of the Amaryllidaceae alkaloid biosynthetic pathway may occur in spatially distinct tissues in *H. papilio*, consistent with the notion of tissue-specific partitioning of specialized metabolism. The clustering of precursors and intermediates was significantly associated with their enrichment in the apical bulb leaf, likely corresponding to the youngest and most biosynthetically active tissue. These results are partially consistent with Mehta et al’s hypothesis that biosynthesis is more active in young tissues of leaves, even though they did not investigate young apical leaves (Mehta *et al*., 2024). However, these compounds were also frequently detected in the epidermis of the bulb mid-section and in the leaf base and mid-section, indicating that early biosynthetic steps may occur across both internal and peripheral tissues. This broader spatial distribution is in line with Haist et al, who reported high alkaloid concentrations in inner bulb tissues, but also observed signal in more superficial tissues, depending on compound type.

The consistent association of haemanthamine with precursors across PCA, t-SNE, and hierarchical clustering further suggests that this compound may be synthesized closer to the precursor zone than previously assumed. In contrast, galanthamine, 11β-hydroxygalanthamine, and papiline-type alkaloids clustered with features such as the root endodermis, leaf tip vascular bundles, and epidermal zones of distal organs, together with the detection of high levels of end-products in bulb and leaves mucilage, this suggests that late-stage tailoring may occur in more specialized cell layers, vascular-associated cells, distinct from the tissues synthesizing early intermediates. These spatial relationships reveal a likely progression from early, distributed biosynthesis toward more compartmentalized localization of end-products. Alternatively, active transport mechanisms could mediate the relocalization of galanthamine from its site of synthesis to specialized storage compartments, potentially minimizing self-toxicity or optimizing resource allocation within the plant. These patterns should be interpreted as exploratory correlations rather than definitive biosynthetic maps, given the qualitative or semi-quantitative nature of the MSI analysis and the sampling that was performed during the vegetative stage. Nonetheless, the consistent separation between precursor and galanthamine-type compounds across unsupervised clustering, PCA, and t-SNE, along with their reproducible association with distinct tissue features, offers a strong foundation for future work that combines enzyme localization, gene expression, and transport assays to test these hypotheses experimentally.

While multivariate analyses have been increasingly adopted in metabolomics datasets, their application to MSI spatial profiles in plants remains limited, particularly for specialized metabolites. The integration of PCA, t-SNE, and hierarchical clustering with MSI datasets allowed the identification of structured tissue-level patterns that would not have been evident through descriptive visualization alone. Recent advances in plant MSI analysis increasingly highlighted the importance of such approaches for uncovering hidden biological organization (Boughton *et al*., 2016; Dong *et al*., 2016; Heyman and Dubery, 2016; Li *et al*., 2024), and our study extends these concepts to the investigation of specialized metabolite biosynthesis and transport in Amaryllidaceae.

In a nutshell, the distribution patterns of AAs observed in the three organs of *H*. *papilio*—leaves, bulbs, and roots—provide valuable insights into the potential biosynthetic pathway arrangement of galanthamine (Figure 8). The data suggest that the early steps of AA biosynthesis are enriched in apical leaves, and the biosynthesis of galanthamine likely occurs in distinct tissue-specific compartments, with varying degrees of metabolic exchange between these regions. In the leaves and bulbs, the pathway appears to be localized primarily in the epidermal tissues and vascular areas, while precursors were comparatively enriched in the apical leaves. The presence of key intermediates in both tissues indicates that the biosynthetic reactions may occur in either compartment, with potential exchange of metabolites between the epidermis and vascular areas. This suggests a dynamic interplay between these regions, where specific steps of the pathway could be restricted to either the epidermis or the vascular tissues, followed by the transport of intermediates to the other compartment for subsequent reactions. In contrast, the roots exhibit a more compartmentalized biosynthetic profile. The limited exchange of intermediates observed in the roots, compared to the leaves and bulbs, implies that the reactions may be more spatially restricted. This compartmentalization could reflect a specialized metabolic strategy in the roots, potentially optimizing the utilization of resources or minimizing the exposure of toxic intermediates to other tissues.

**Figure 8.**
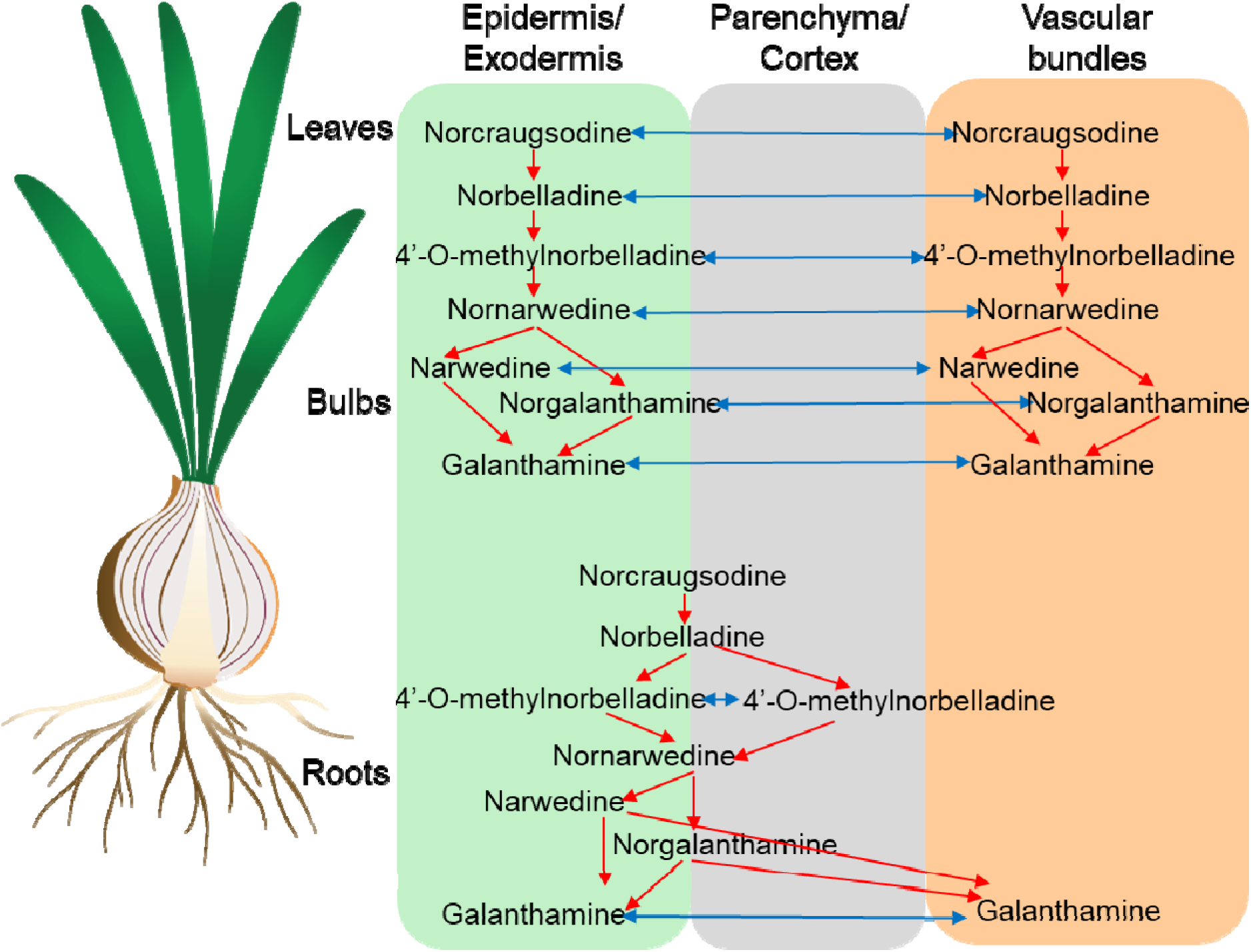
The proposed biosynthetic organization of galanthamine in *Hippeastrum papilio* is illustrated, with red arrows denoting biochemical steps and blue arrows indicating potential alkaloid transportation. The pathway in the top is restricted to the leaves and the bulbs and the bottom pathway belongs to the roots.

It was shown that the change in withanolides in the aeroponically grown medicinal plant *Withania somnifera,* where withanolides usually accumulate in roots, starts to accumulate in the leaves when they grow aeroponically (Xu *et al*., 2011). Thus, differences in growth conditions complicate the comparison of the biosynthesis of AA of *H*. *papilio* with geophytic species like *N*. *tazetta* or *Narcissus* cv. Tête-à-Tête (Mehta *et al*., 2024). Based on our observations, the biosynthesis of AA in *H. papilio* does not exhibit an organ-specific pattern. This idea is supported by the detection of biosynthetic gene expression in all the organs, although the level of expression varies. For example, *norbelladine synthase* and *norcraugsodine/noroxomaritidine reductase* and *4’-O-methyltransferase* involved in the formation of precursors and intermediates are detected in all organs of *Leucojum aestivum* and *Narcissus papyraceus* (Koirala *et al*., 2024; Majhi *et al*., 2023). Similarly, the transcripts of N-methyltransferase, which can be involved in the formation of galanthamine, 3’-O,N-dimethylnorbelladine, 3,4’-O,N-trimethylnordelladine were present in all the studied organs of *L*. *aestivum*, *Lycoris radiata*, and *H*. *papilio* (Liyanage *et al*.). Some steps may be tissue-specific; however, further investigation, including multi-omics or single-cell-level analysis, is needed to determine the exact information that elucidates the pathway.

### 4.5. New metabolites uncovered in *H. papilio*

By leveraging multiple databases integrated with Metaspace, we identified several previously unreported AAs in *H*. *papilio*, expanding the known phytochemical profile of this species. Notably, our findings also provide evidence for compounds associated with colchicine biosynthesis, a pathway previously documented only in *Lycoris radiata* (Yan *et al*., 2018). The detection of multiple intermediates in this pathway supports the possibility of shared or convergent biosynthetic routes within the Amaryllidaceae family. Additionally, we observed diverse alkaloid groups beyond the typical Amaryllidaceae-type structures, underscoring the metabolic complexity of this plant family (Berkov *et al*., 2020; Jin and Yao, 2019). Beyond targeted alkaloid profiling, this non-targeted approach proves valuable for broader metabolomic investigations. The simultaneous detection of flavonoids, lignans, pigments, and other specialized metabolites highlights the potential of MALDI-MS in studying metabolic correlations, plant-environment interactions, and biosynthetic networks. Such comprehensive analyses could pave the way for future research on the ecological and physiological roles of diverse metabolite classes in plants.

In summary, this study employed MALDI-MSI to investigate the spatial distribution of AAs in *H*. *papilio* leaf, bulb, and root tissues. Galanthamine accumulated particularly in epidermal and vascular regions. Leaves showed consistent alkaloid distribution across ages and positions, while bulbs exhibited variations between outer and inner scales. Roots displayed compartmentalized alkaloid distribution, with higher concentrations at the proximal end. Biosynthetic intermediates and minor alkaloids were present, albeit at lower intensities, and were specifically concentrated in the bulb apical leaves. The findings highlight tissue-specific alkaloid distribution patterns, suggesting specialized biochemical pathways in different plant regions.

These findings contribute to a deeper understanding of the ecological and physiological significance of alkaloids in *H*. *papilio* and highlight the importance of tissue-specific metabolic profiling in plant research. Future studies could explore the subcellular localization of these alkaloids and their biosynthetic pathways to further elucidate their functional roles and regulatory mechanisms in *H*. *papilio*. Additionally, comparative studies across different plant species with varying ecological niches could provide broader insights into the adaptive significance of alkaloid distribution patterns.

## 5. Conclusion

To the best of our knowledge, this is the first comprehensive study of the spatial metabolic arrangement of alkaloids in an Amaryllidaceae species. Our analysis revealed that galanthamine-type and papiline-type AAs exhibited a consistent spatial arrangement in the organs of *H*. *papilio, whic*h differed from that of intermediates and haemanthamine. The abundance of AAs in epidermal and vascular tissues highlights their potential role in defense mechanisms, while the presence of biosynthetic intermediates suggests widespread alkaloid biosynthesis. In leaves, the spatial distribution of metabolites appears interdependent: areas with early intermediates consistently show downstream metabolites, and conversely, regions with lower detection of precursors demonstrate reduced detection of AAs. In bulbs, while intermediates and end-products were detected in older scales and basal plates, intermediates were more specific to the apical leaves. In the basal plates, most localized in vascular bundles, while galanthamine was also abundant in the parenchyma. In roots, AAs were less prevalent in the vascular bundles than in the cortex, with galanthamine and its derivative being a notable exception. This unique accumulation pattern of galanthamine potentially indicates distinct transport or storage mechanisms compared to other alkaloids. These findings provide crucial insights into the tissue-specific biosynthesis and localization of AAs, emphasizing the coordinated nature of their production and storage across various plant organs. This knowledge advances our understanding of the metabolic architecture of *H*. *papilio* and sets the stage for exploring its potential biotechnological applications.

## Author Contribution

**Nuwan Sameera Liyanage:** Conceptualization, Methodology, Experiments, Analysis, Writing-Original draft, Writing-Reviewing and Editing; **Natacha Merindol:** Analysis, supervision, Writing-Reviewing and Editing; **Kenneth Munk Pedersen:** LC-MS and analysis, Writing-Reviewing and Editing; **Christian Janfelt:** Conceptualization, Methodology, Resources, supervision, funding acquisition, Writing-Original draft, Writing-Reviewing and Editing **Isabel Desgagné-Penix:** Conceptualization, Resources, supervision, funding acquisition, Writing-Original draft, Writing-Reviewing and Editing.

## Acknowledgements

The authors thank Dr. Marcus Daniel Brandbjerg Bohn Lorensen (University of Copenhagen) and Rohith Grandhi (Université du Québec à Trois-Rivières) for their valuable input and contributions. We also thank Prof. Antonio Evidente (Universitario Monte Sant’Angelo, Naples, Italy) for providing some alkaloid standards. During the preparation of this work, the authors used ChatGPT version 4.0, a free AI language model, in order to write the scripts in Python, correct grammatical errors and enhance readability. After using this tool, the authors reviewed and edited the content as needed and take full responsibility for the content of the publication.

## Data availability

Raw data in the form of imzML files of the presented data has been uploaded to the Metaspace platform (https://metaspace2020.org/project/Hippeastrum_paplio_MALDI) and annotated against multiple publicly available databases. Access to the reviewers by https://metaspace2020.org/api_auth/review?prj=0247bb56-4f1e-11ef-86c3-fbc99e2b8fbc&token=_Xv46BbxCZf8

---

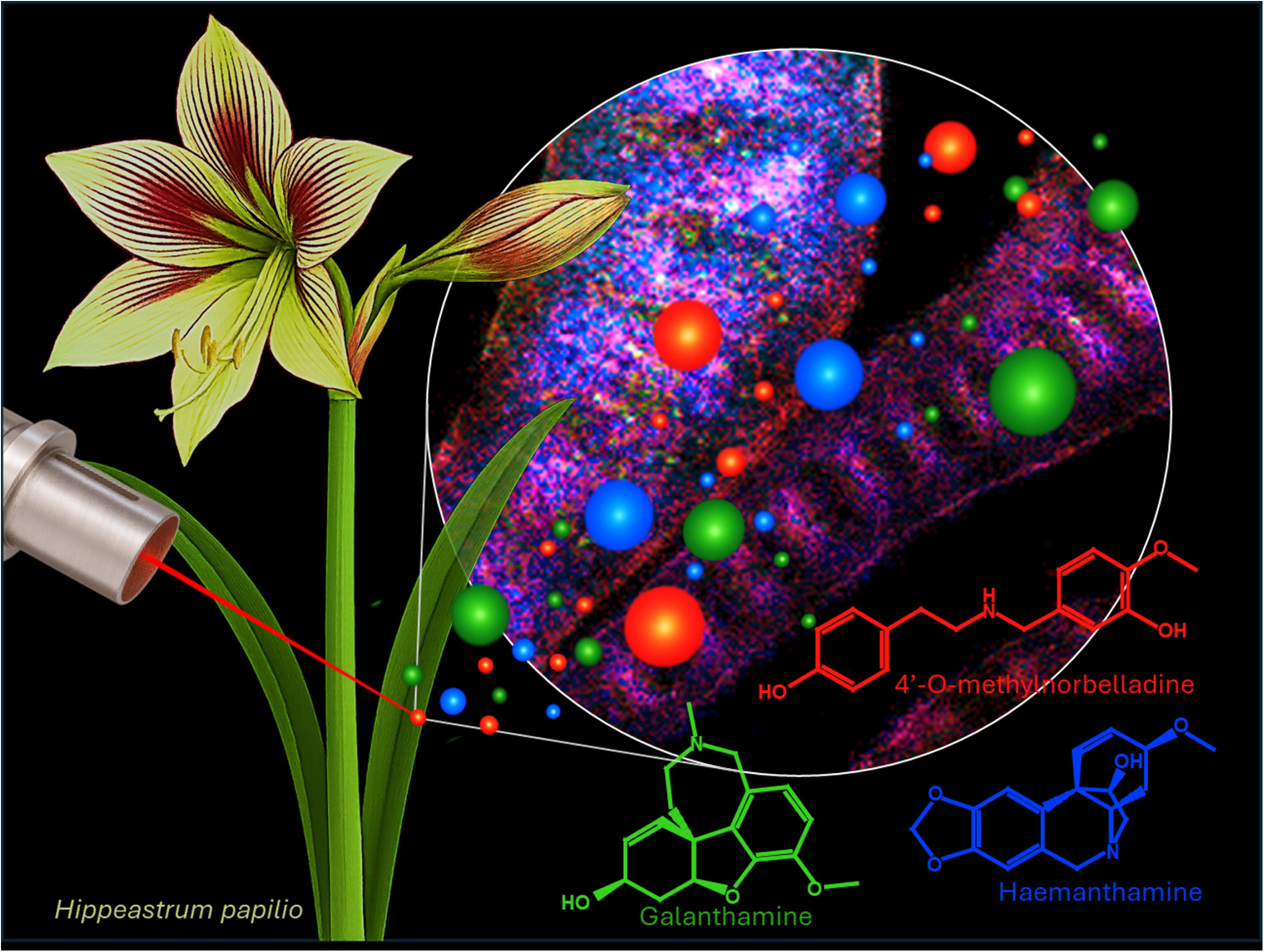

